# Cross-species analysis identifies genotype-driven vulnerabilities in lung adenocarcinoma

**DOI:** 10.1101/2025.08.24.671070

**Authors:** Angeliki Karatza, Maayan Pour, Jaylen M Powell, Yuan Hao, Dimitris Nasias, Yi Jer Tan, Alfonso Lopez, Metamia Ciampricotti, Chengwei Peng, Alvaro Quintanal Villalonga, Yeuan Ting Lee, Ruveyda Ayasun, Muhammad Elnaggar, Priyanka Sahu, Igor Dolgalev, Hai Hu, Luis Chiriboga, Cynthia Loomis, Daniel Andrussier, Magdalena Ploszaj, Aristotelis Tsirigos, Peter Meyn, Fred P Davis, Nicolas Stransky, Gromoslaw A Smolen, John Thomas Poirier, Jonathan So, Mark Philips, Harvey Pass, Dafna Bar Sagi, Itai Yanai, Kwok-Kin Wong

**Author notes:** **Corresponding Authors** Kwok-Kin Wong (K.K.W.), and Itai Yanai (I.Y.). Contributed equally (Angeliki Karatza, Maayan Pour). Contributed equally (Jaylen M Powell, Yuan Hao).

## Abstract

While three major genetic alteration subsets, characterized by mutations in *STK11, KRAS*, and *EGFR*, are seminal in driving tumorigenesis in LUAD, their distinct effects on tumor cells and the tumor microenvironment are not fully understood. Here, we map critical oncogenic subset-specific vulnerabilities by identifying conserved cell-type-specific reprogrammings between human and mouse LUAD. Through harmonized scRNA-seq analysis of 57 human and 18 mouse specimens, we unveil that genetic alterations impose genotype-specific immune imprints on the tumor microenvironment: KRAS is associated with a transitional immune state, whereas STK11 and EGFR mutations define discrete and contrasting immune phenotypes. We find that STK11-mutant tumors exhibit complement and interferon-rich immune microenvironments while EGFR-mutant tumors harbor a naive T cell-rich phenotype accompanied by a global HLA downregulation, stress-responsive alveolar macrophages marked by MARCO. In the epithelial compartments, cross-species analysis reveals metabolic dependencies, including *OGDH* in EGFR- and *PDE4D* in STK11-mutant cells.

**Statement of Significance:** This study delineates oncogenotype-specific molecular and immune imprints in lung adenocarcinoma, revealing STK11-, KRAS-, and EGFR-mutated tumors as transcriptionally and immunologically discrete ecosystems. By mapping epithelial vulnerabilities alongside immune imprints, we unveil genotype-imposed dependencies that inform the rational development of precision therapies with direct translational relevance.

## Introduction

Lung cancer remains the leading cause of cancer-related deaths, accounting for over 20% of total cancer mortality in the United States (1). Non-small cell lung cancer (NSCLC) accounts for 85% of lung cancer cases, of which lung adenocarcinoma (LUAD) constitutes 40% (2). The standard of care for LUAD patients depends on the stage of the disease at diagnosis, and it typically encompasses surgical intervention, chemotherapy, and radiation therapy. Over the past few decades, integrating immunotherapy and targeted therapies into the treatment paradigm has significantly improved the clinical efficacy and survival rates of patients with LUAD (3).

Advances in next-generation sequencing (NGS) technologies have greatly enhanced the detection of oncogenic alterations, biomarker discovery, and patient stratification for targeted therapies and immunotherapies (4). For instance, patients with LUAD harboring *EGFR*-activating mutations have benefited from therapies with tyrosine kinase inhibitors (TKIs), which disrupt the abnormal *EGFR* signaling pathway (5). Similarly, in LUAD patients with oncogenic *KRAS*-activating mutations, the development of KRAS-specific inhibitors and degraders has significantly transformed the treatment landscape for this patient subset (6), (7).

Despite the clinical efficacy of targeted therapies, the lack of response in many patients and the emergence of drug resistance present notable challenges. Consequently, combining targeted therapies with other therapeutic agents has shown promise in improving response rates and mitigating drug resistance, thus offering a potential strategy for enhancing treatment efficacy. For example, the combination of targeted therapies with immune checkpoint inhibitors such as programmed cell death protein 1 (PD-1) and cytotoxic T-lymphocyte-associated protein 4 (CTLA-4) has significantly improved clinical outcomes in patients with LUAD (8),(9). Additionally, targeted therapies combined with metabolic inhibitors, such as those targeting glutamine and mitochondrial metabolism, are currently undergoing clinical trials (10). These examples underscore the urgent need to identify novel therapeutic targets within tumor cells and the tumor microenvironment (TME) to advance therapeutic strategies.

Numerous studies have profiled human LUAD by single-cell RNA sequencing (scRNA-seq). In a recent study, researchers combined multi-parameter mass cytometry with scRNA-seq to identify cancer-specific rather than organ-specific gene signatures in the immune compartment of LUAD patients (11). Other studies have employed scRNA-seq to compare human LUAD with other histological lung cancer subtypes, such as lung squamous carcinoma (Wu, 2021 #74}. Similarly, single-cell methodologies have been employed to characterize the dynamic changes in lung adenocarcinoma during treatment (12).

Although scRNA atlases have substantially enhanced our understanding of the LUAD landscape and heterogeneity, several questions remain unresolved. Previous studies have suggested that oncogenic driver mutations play a crucial role in shaping the TME composition. For example, the prevalence of neutrophil-rich *milieu* in *KRAS; STK11*-mutated lung adenocarcinoma tumors has been reported (13). A lack of T cells and high levels of T-cell exhaustion and immune suppression have also been reported in this subset of LUAD (14), (15). Thus, the study of the potential dependence of TME heterogeneity on specific oncogenic drivers at the single-cell level could reveal the role of the TME in tumor progression and response to treatment.

One of the main constraints is the limited genomic information available for freshly resected lung tumor specimens, which are typically processed for scRNA-seq analyses. In this study, we utilized single-nucleus RNA sequencing (snRNA-seq) and scRNA-seq to examine the tumor and stromal compartments across major oncogenic driver subsets in LUAD. We conducted a comprehensive cross-species single-cell transcriptomic comparison of three distinct LUAD subgroups: *STK11*, *KRAS*, and *EGFR*.

For human analyses, we generated a snRNA-seq dataset from frozen naïve LUAD specimens, enabling genotypic characterization and stratification. For our murine analyses, we used three well-characterized genetically engineered mouse models (GEMMs) of lung adenocarcinoma, each designed to recapitulate one of the three major genetic alterations observed in human LUAD. These GEMMs are invaluable tools in lung cancer research, as they closely mimic human lung cancer pathogenesis by introducing specific genetic alterations into the mouse genome. Utilizing lung cancer GEMMs allows the observation of lung tumor progression and development in a physiologically relevant context (16), (17). In contrast, while human samples, patient-derived xenografts (PDXs), and human cell lines accurately recapitulate tumor heterogeneity, they are unsuitable for immune system and immunotherapy studies because of the lack of intact immune systems. GEMMs, with their preserved immune systems, offer superior platforms for investigating immune responses and testing immunotherapy. Furthermore, these GEMMs are instrumental in the preclinical testing of new therapeutics, allowing for the assessment of drug efficacy and investigation of resistance mechanisms.

In this study, we provide a holistic survey revealing that the genetic underpinnings driving LUAD not only govern the transcriptional circuitry of malignant cells but also profoundly reprogram the tumor microenvironment by reshaping the cellular *milieu* and inducing mutation-specific transcriptional states across immune and stromal compartments.

Deciphering these mutation-driven functional states holds significant clinical promise, offering a path toward more precise therapeutic stratification, particularly in immunotherapy and targeted therapy, and ultimately enhancing patient outcomes through tailored intervention strategies.

## Results

### Convergent cross-species analysis as a platform to delineate mutation-driven remodeling of cellular and transcriptional landscapes in LUAD

To assess mutation-driven heterogeneity in human LUAD, we generated a snRNA-seq dataset using 57 frozen resections from 55 primary, treatment-naïve patients (replicates were obtained for two patients). In addition to snRNA-seq, each sample underwent bulk DNA whole-exome sequencing (**Fig. 1A, 1B,** and **Supplementary Fig. S1A**). Based on whole-exome data, samples were stratified into three subsets according to the oncogenic driver mutations they harbored: *STK11, KRAS,* and *EGFR* (**Fig. 1A, 1C, and Supplementary Fig. S1A and S1B**). We then identified the major cell types using cell-type-specific gene markers in the single-nucleus data (**Fig. 1D, 1E,** and **Supplementary Fig. S1C**). Neutrophils were not detected, as expected from the snRNA-seq dataset (18). Epithelial (malignant) cell transcriptomes clustered highly by patient, while endothelial cells, fibroblasts, macrophages, T/NK cells, and B cells formed cell-type-driven clusters **(Fig. 1B and 1D**). We examined the intra- and inter-genotype cell type composition and found that epithelial cells were the most prevalent in the snRNA-seq dataset **Fig. 1F and Supplementary Fig. S2A**).

**Figure 1.**
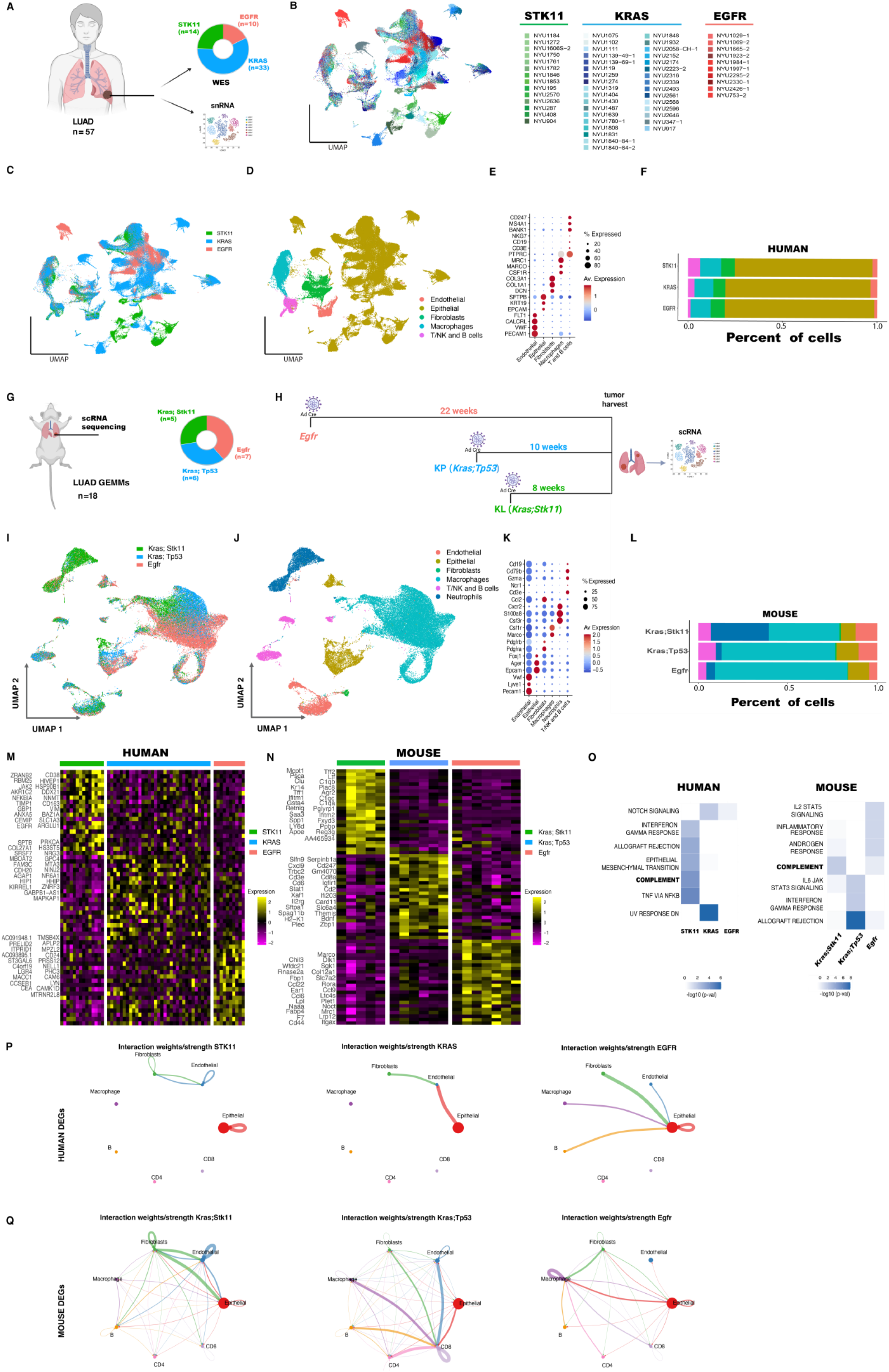
Convergent cross-species genotype analysis identifies the complement pathway as a hallmark of STK11*-*mutated LUAD. **A**) Schematic illustration of the pipeline and distribution of the three major genotype subsets in human lung adenocarcinoma samples. **B–D**) UMAP visualization of snRNA-seq data from 57 human lung adenocarcinoma specimens by patient ID **(B),** genotype of the oncogenic driver **(C),** and cell type **(D)**. **E**) Dot plot displaying the average expression levels of gene markers characteristic of cell-type clusters in human lung adenocarcinoma. **F)** Stacked bar plots of cell-type percentage stratified by genotype in human LUAD. **G**) Pipeline and distribution of the three major genotype subsets among the mouse lung adenocarcinoma samples. **H**) Schematic depiction of genetically engineered lung adenocarcinoma mouse models used in this study, with the timeline of tumor induction. **I-J**) The UMAP projection of scRNA-seq data stratifies mouse lung adenocarcinoma samples by oncogenic mutation **(I)** and cell type **(J**). **K**) Dot plots demonstrate the average expression levels of gene markers, allowing the identification of cell type clusters in mouse lung adenocarcinoma. **L)** Stacked bar plots of cell-type percentage stratified by genotype in mouse LUAD GEMMs. **M, N**) Heatmap displaying the top DEGs of the global collective analysis of cancer and TME cells averaged by patient **(M)** or mouse **(N)** samples among the three LUAD genotypes. **O**) Pathway enrichment analyses of human and mouse DEGs identified the complement pathway as an STK11 global signature. **P**) Cell-cell communication analysis of the DEGs among the three human LUAD genotypes. **Q**) Cell-cell communication analysis of the DEGs among the three LUAD GEMMs.

To ensure translational relevance and cross-species robustness, we implemented a comparative mouse-human analytic framework, enabling the identification of conserved and biologically salient transcriptional programs across species. Specifically, we employed three genetically engineered mouse models (GEMMs) of lung adenocarcinoma (LUAD)—Kras; Stk11, Kras; Tp53, and Egfr—each of which mirrors a clinically relevant oncogenic driver mutation (**Fig. 1G**). To enhance experimental fidelity and reproducibility, three independent experimental cohorts were established, each comprising multiple biological replicates (**Supplementary Fig. S3A–C**). Tumors were harvested synchronously at defined time points post-Cre induction, corresponding to uniform stages of tumor progression, as delineated in the mouse experimental schema **(Fig. 1H)**. Importantly, single-cell RNA sequencing (scRNA-seq) was performed on each individual tumor to capture intra-tumoral heterogeneity at high resolution.

For each GEMM, single-cell transcriptomic datasets from the three independent experiments were integrated for downstream comparative analyses (**Supplementary Fig. S3D–F**). Consistent with our human LUAD analyses, we performed cell-type annotation using canonical lineage markers to classify major tumor and stromal compartments (**Fig. 1I–K**). STK11-deficient tumors exhibited a pronounced enrichment of neutrophils **(Fig. 1L and Supplementary Fig. S2B**), consistent with prior findings (13), thereby validating the fidelity of the GEMMs in recapitulating genotype-specific microenvironmental features.

Previous bulk RNA-seq studies of human patients’ tumors have highlighted immunosuppressive phenotypes in STK11-mutant LUAD (19). Extending this work, we performed pseudo-bulk differential expression analyses across genotypes, where we used our snRNA-seq to average the expression profiles per cell type for each tumor, to avoid biases based on cell type abundance differences. We delineated genotype-specific differentially expressed genes (DEGs) across KRAS, STK11, and EGFR mutation groups (**Fig. 1M**). Notably, STK11-mutant tumors exhibited upregulation of previously implicated therapeutic targets such as *HSP90B1* (19), *CD38* (20), and *AKR1C2*(21). Conversely, *LYN* (22), *LRG4* (23), and *CD24* (24) were among the most prominent DEGs in EGFR-mutant tumors, consistent with prior reports.

To benchmark these human findings in a preclinical context, we applied analogous DEG analyses to the scRNA-seq data from the mouse models. Following cell-type-specific averaging and exclusion of neutrophils (to align with human datasets), we derived a cross-species DEG matrix (**Fig. 1N**). While direct overlap in DEG identity between species was limited, gene set enrichment analyses uncovered significant convergence at the pathway level. Notably, both human and mouse STK11-deficient tumors demonstrated enrichment of the complement cascade, implicating this pathway as a conserved hallmark of STK11 loss (**Fig. 1O**).

Subsequently, we constructed the cell-cell interactome in both human **(Fig. 1P and Supplementary Fig. S2C)** and mouse **(Fig. 1Q and Supplementary Fig. S2D)** LUAD using differentially expressed ligand-receptor gene pairs. In the mouse models, the Kras; Stk11 subset exhibited prominent fibroblast-endothelial signaling, the Kras; Tp53 subset showed enrichment in CD8⁺ T cell-mediated interactions, and the Egfr subset was characterized by macrophage– macrophage communication. Cross-species analyses revealed conserved stromal circuits in the STK11-deficient subset, including fibroblast–fibroblast, endothelial–endothelial, and fibroblast– endothelial interactions. Additionally, conserved intra-epithelial communication was enriched in both STK11- and EGFR-mutant subsets.

In summary, our integrative, cross-species single-cell analyses uncover genotype-specific intercellular signaling architectures in LUAD, with the complement pathway and conserved stromal and epithelial interactomes emerging as hallmarks of STK11 deficiency. These findings highlight the importance of genotype-driven cellular dependencies and nominate context-specific cell-cell interactions as potential therapeutic entry points across distinct molecular LUAD subtypes.

### Genotype-instructed Cell-Cell Communication networks in lung adenocarcinoma

To dissect the landscape of genotype-specific cell-cell communication in LUAD, we conducted a comprehensive interactome analysis across the three major oncogenic drivers. Although the number of cell-cell interactions by genotype analyses in human LUAD did not yield striking differences **(Fig. S4A),** we quantified the proportion of ligand-receptor interactions attributed to key signaling pathways by genotype in human LUAD **(Fig. 2A)**. Strikingly, this analysis uncovered a pronounced depletion of MHC-I and SIRP module interactions in EGFR-mutant tumors, suggesting a potential evasion of antigen presentation and phagocytic clearance in this oncogenic context. Furthermore, the CD46 interactome, central to complement cascade modulation, was selectively absent in EGFR-mutant tumors, highlighting a genotype-specific immune regulatory deficit. Distinct genotype-specific pathway interactomes were uncovered, with STK11 tumors enriched for the TGFb pathway, KRAS tumors for the GALECTIN axis, and EGFR tumors for the GDF signaling network **(Fig. 2B)**. Further dissection of genotype-specific interactomes revealed that TGFb signaling in STK11-mutant tumors is orchestrated by Tgfb1+ endothelial cells, and Lgals9 macrophages mediate GALECTIN signaling. GDF signaling is driven by epithelial cells expressing GDF15 **(Supplementary Fig. S4B).** Galectin-9, a ligand for PD-1 and TIM-3, contributes to T cell exhaustion and has emerged as a candidate for immune checkpoint modulation(25). GDF15, a stress-responsive cytokine, is induced by cellular stress, inflammation, and metabolic dysfunction (26). Next, we validated these genotype-specific signaling pathways in LUAD GEMM subsets. Notably, TGFb signaling exhibited increased cell-cell interactions in the Kras; Stk11 model, while the Galectin pathway was preferentially enriched in the Kras; Tp53 subset. Interestingly, GDF signaling was more prominent in the Kras; Tp53 group **(Fig. 2C and Supplementary Fig. S4C)**. We next investigated genotype-specific epithelial interactomes in human LUAD. EGFR-mutant tumors exhibited a marked absence of both HLA- and CD46-mediated interactions, consistent with MHC class I downregulation and the global loss of CD46 signaling observed in this subset. In contrast, NRXN1-centered interactions were enriched in STK11- and KRAS-mutant tumors, but absent in EGFR, providing a therapeutic opportunity in these subsets of LUAD. Further dissection revealed distinct genotype-specific epithelial ligands: STK11 tumors were characterized by enrichment of LPAR1, FLRT3, and KITLG; KRAS tumors by LAMC2 and LAMB3; and EGFR tumors by TNC, GDF15, and WNT5B. **(Fig. 2D)**

**Figure 2.**
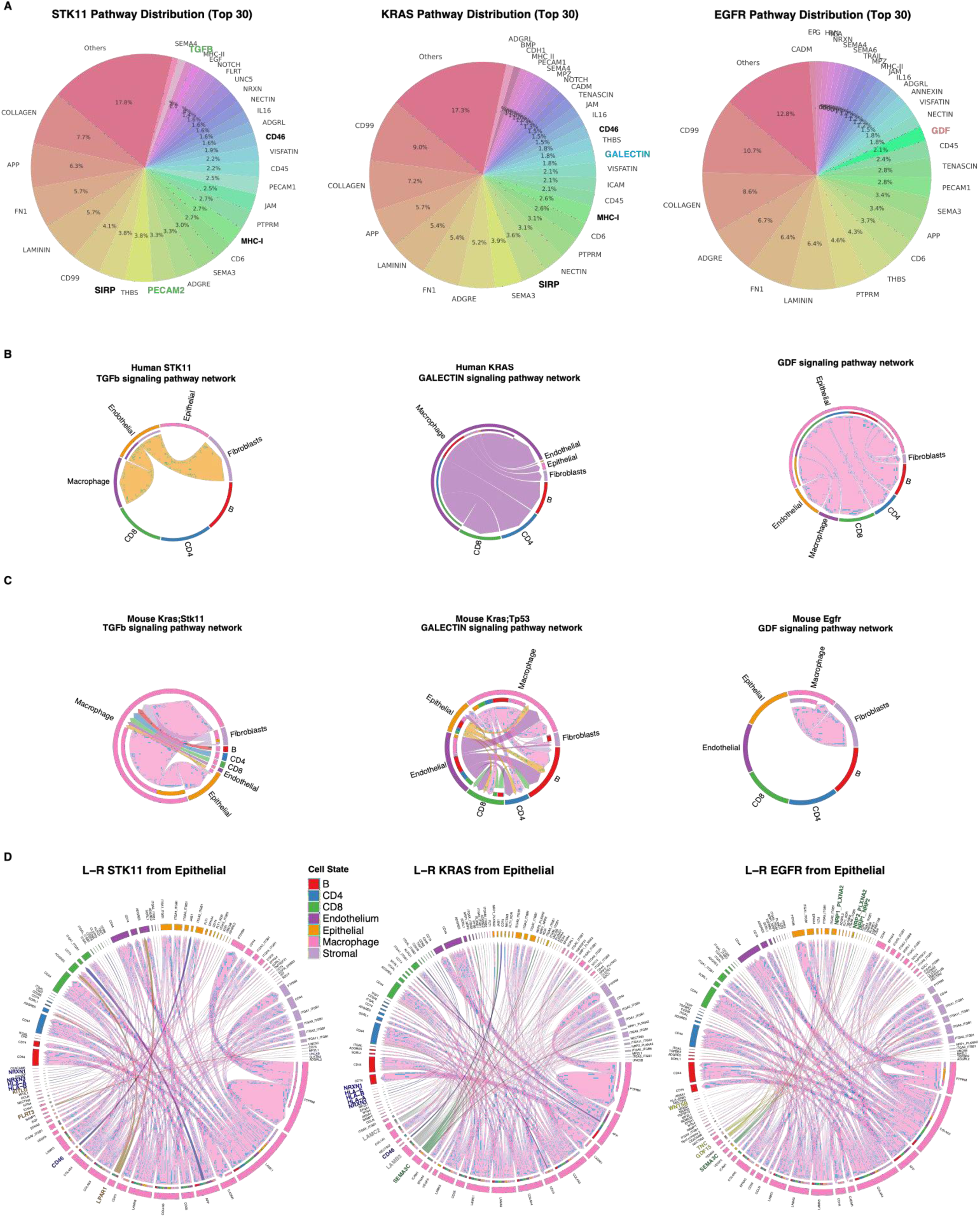
Genotype-instructed cell-cell communication networks in lung adenocarcinoma. **A)** Pie charts visualizing the representation of cell–cell communication pathways across oncogenic genotypes in human LUAD. **B)** Genotype-specific pathway-level interactions in the human LUAD. **C)** Genotype-enriched pathway cell interactions in mouse LUAD. **D)** Circular plots depicting epithelial cell communication networks stratified by genotype in human LUAD, highlighting distinct epithelial interactomes across oncogenic contexts.

Collectively, these cell-cell communication analyses unveil novel genotype-specific epithelial interactomes that define the tumors’ signaling equilibrium and expose putative therapeutic vulnerabilities across LUAD subsets.

### Oncogenotype-driven immune imprints govern the cellular architecture and transcriptional programs of the LUAD tumor immune microenvironment

We employed SCENIC-based regulatory network inference across distinct cellular subsets to interrogate the impact of oncogenic mutations on the transcriptional architecture of immune compartments in LUAD. Hierarchical clustering of significantly enriched regulons in both human **(Fig. 3A and Supplementary Fig. S5A)** and mouse **(Fig. 3C and Supplementary Fig. S5B)** datasets revealed that transcriptional control is predominantly dictated by cellular lineage rather than mutational genotype. Notably, in human LUAD, EGFR-mutant lymphocytes—including both T cells and B cells—exhibited convergent transcriptional regulation, whereas other immune and stromal cell populations were governed by cell–type–specific regulatory programs, largely independent of oncogenic context. Transcriptional regulons that govern the EGFR lymphoid cells include known immune cell transcriptional factors such as NFKB2, and naïve cell markers such as TCF7 (27) **(Fig. 3B)**.

**Figure 3.**
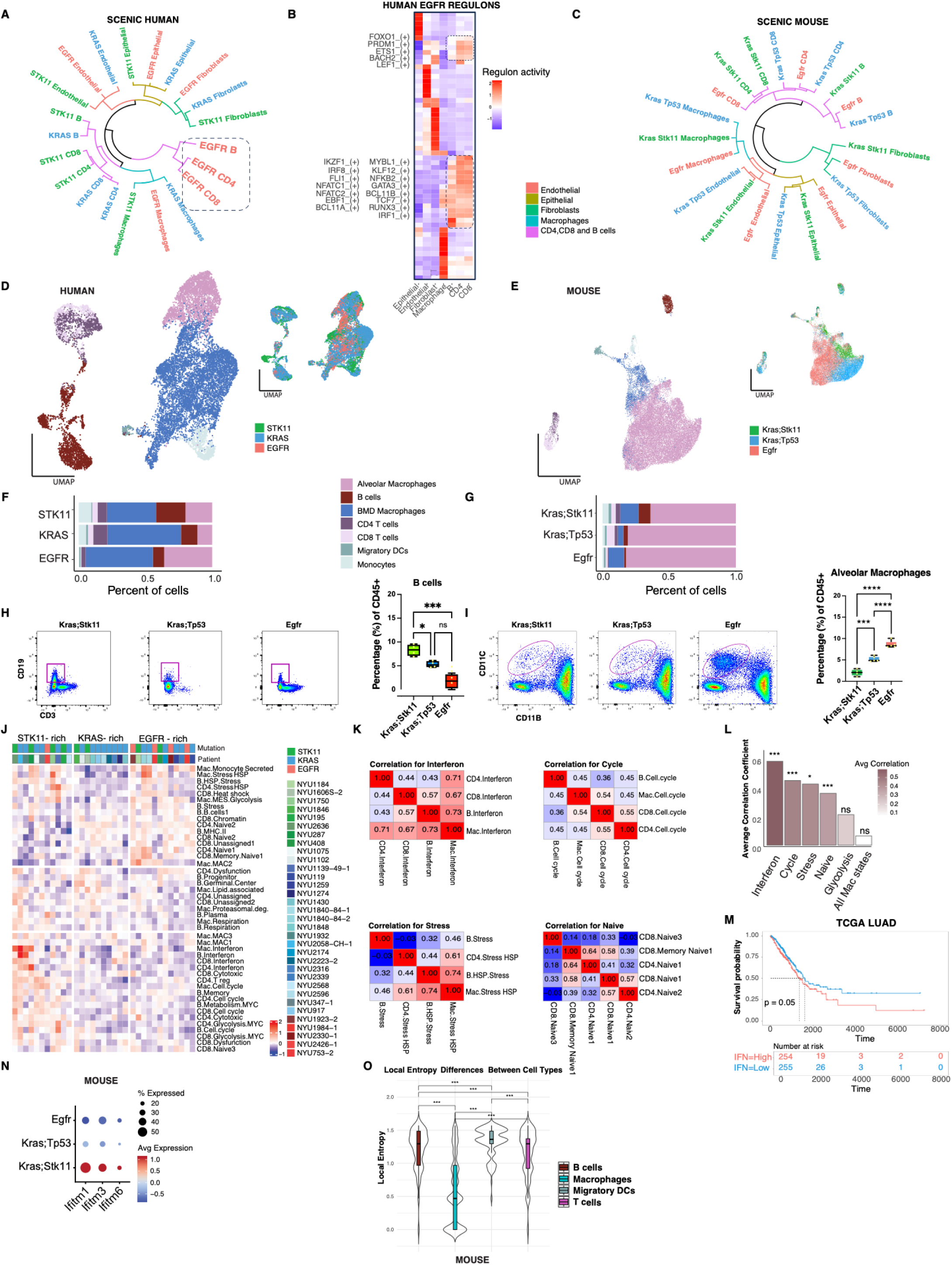
Oncogenotype-driven immune imprints govern the cellular architecture and transcriptional programs of the LUAD tumor microenvironment. **A,C)** Circular dendrograms displaying hierarchical clustering of SCENIC-inferred regulons across human **(A)** and mouse **(C)** LUAD immune cell types. **B)** Heatmap displaying the human transcriptional regulons controlling the cell fate of EGFR lymphocytes. **D, E)** UMAP projections of immune cell populations in human **(D)** and mouse **(E)** LUAD. **F, G)** Stacked bar plots quantifying immune cell composition as a percentage of total immune cells in human **(F)** and mouse **(G)** LUAD. **H, I)** Flow cytometry analysis comparing B cell **(H)** and alveolar macrophage **(I)** populations across LUAD GEMMs stratified by genotype. **J)** Heatmap of average scores of immune-related gene signatures in the human LUAD tumor immune microenvironment. **K)** Correlation heatmaps of immune signatures of Interferon (top left), Stress (bottom left), Cell cycle (top right) and Naive (bottom right) cellular programs from different immune cell types, for their score across human LUAD patients. **L)** Average correlation coefficients of the immune gene signatures within the LUAD tumor microenvironment, that presented in K. **M)** Kaplan–Meier survival analysis in the TCGA LUAD cohort, stratified by high versus low expression of the interferon gene signature. **N)** Dot plot of average expression and percent of cells expressing interferon program-related genes in mouse LUAD GEMMs. **O)** Entropy analysis of immune cell diversity of drive mutation in mouse GEMMs. Lower entropy suggests higher order by drive mutation.

Recognizing that oncogenic lesions may shape the broader immune *milieu*, we next deconstructed the LUAD tumor immune microenvironment (TIME) in a genotype-stratified framework. Unsupervised UMAP embedding delineated canonical immune cell populations in both human **(Fig. 3D)** and mouse **(Fig. 3E)** TIME. Strikingly, comparative analysis across major oncogenic drivers unveiled a consistent enrichment of B lymphocytes in STK11-mutant tumors, in contrast to the preferential accumulation of alveolar macrophages in EGFR-mutant LUAD both in human and mouse **(Figs. 3F and 3G)**. These compositional biases were functionally validated *in vivo* using flow cytometric profiling of LUAD GEMMs, which confirmed the selective expansion of B cells in STK11-driven tumors **(Fig. 3H)** and a marked increase in CD11c⁺F4/80⁺ alveolar macrophages in EGFR-driven models **(Fig. 3I)**.

Next, to further dissect the immunogenomic architecture of the LUAD tumor immune microenvironment, we computed the average expression of curated gene modules, reflecting distinct functional states across key immune populations—macrophages, B cells, CD4⁺, and CD8⁺ T cells—based on transcriptional programs previously defined (28). Hierarchical clustering of these immune state profiles across patients delineated three transcriptionally and genetically distinct TIME imprints **(Fig. 3J)**. Notably, one cluster was markedly enriched for EGFR-mutant tumors, exhibiting elevated expression of naïve T cells and stress-related programs across multiple immune compartments, consistent with a blunted immunogenic state. In line with this, our cell– cell communication analyses identified GDF15, a cytokine induced by cellular stress, as a key EGFR-specific signal. The naïve T cell skewing aligns with the observed downregulation of MHC class I machinery, suggesting impaired antigen presentation as a mechanism of immune evasion. In contrast, a second cluster, dominated by STK11-mutant tumors, was characterized by uniformly high interferon and cell cycle signaling activity across macrophages and lymphocytes, while the third cluster, largely composed of KRAS-driven tumors, displayed intermediate features.

Analyses of the non-immune tumor microenvironment did not recapitulate the same clustering among endothelial cells and fibroblasts **(Supplementary Figs. S5C and S5D**); however, STK11-mutant tumors were enriched for interferon-responsive genes within both compartments (Supplementary Fig. S5E). Conversely, consistent with prior findings, EGFR-mutant tumors exhibited downregulation of MHC class I genes **(Supplementary Fig. S5F).**

Correlative analysis across immune subsets revealed a positive correlation between the aforementioned TIME programs (interferon, cycle, stress, and naïve) **(Fig. 3K)**. Furthermore, stress programs were significantly positively correlated, particularly in EGFR-mutant tumors, where they co-expressed with naïve T cell signatures, suggesting a state of chronic inflammation coupled with ineffective immune priming. This stress-enriched phenotype in EGFR-driven TIME was especially pronounced in macrophages, aligning with transcriptional hallmarks of M2-like polarization and chronic inflammatory adaptation. Furthermore, the T cell naïve observation aligns with the enrichment of TCF7-associated naïve transcriptional programs identified by SCENIC analysis in EGFR-mutant T cells. **(Fig. 3L)**.

Extending these insights to clinical outcomes, integrative survival analysis of the TCGA-LUAD cohort revealed that heightened expression of the interferon-related signature across immune cells was significantly associated with poor prognosis, underscoring the pathogenic relevance of this transcriptional axis in STK11-mutant lung adenocarcinoma **(Fig. 3M)**. In murine models, we further validated the interferon-high phenotype by showing that Ifitm gene family members were selectively upregulated in the Kras; Stk11 compound mutant TIME, mirroring the human data **(Fig. 3N)**.

As the same function-gene signatures from different TIME cell types showed high similarity in their levels in patients, we aimed to test whether differences exist in the genotype-driven phenotypes between these cell types. We used entropy analysis on the mouse immune cells and unveiled a pronounced stratification of macrophages according to oncogenic mutational status **(Fig. 3O)**.

Comprehensive analyses of the LUAD tumor immune microenvironment revealed that oncogenic alterations not only reshape the cellular composition of the immune compartment but also profoundly reprogram its transcriptional landscape. Collectively, these findings define mutation-aligned immune imprints—a stress-naïve EGFR-rich TIME, a cycling interferon-inflamed STK11-rich TIME, and an intermediate KRAS TIME—each representing a distinct immunological ecosystem with translational relevance. Furthermore, among the diverse immune populations, macrophages exhibited the most pronounced transcriptional and phenotypic stratification in response to oncogenic mutations.

### Genotype-guided profiling of LUAD macrophages reveals *MARCO* and *C1Q* as genotype-specific macrophage genes

Building upon our findings on genotype-dependent macrophage gene signature expression, we sought to further investigate genotype-driven genes within the myelomonocytic compartment with therapeutic potential. To identify specific genes and uncover novel genotype-derived therapeutic targets, we conducted a comprehensive global comparison of macrophages at the single-cell level, identifying differentially expressed genes (DEGs) across the three oncogenic subsets in both human **(Fig. 4A and Supplementary Table S1)** and mouse **(Fig. 4B and Supplementary Table S2)** macrophages.

**Figure 4.**
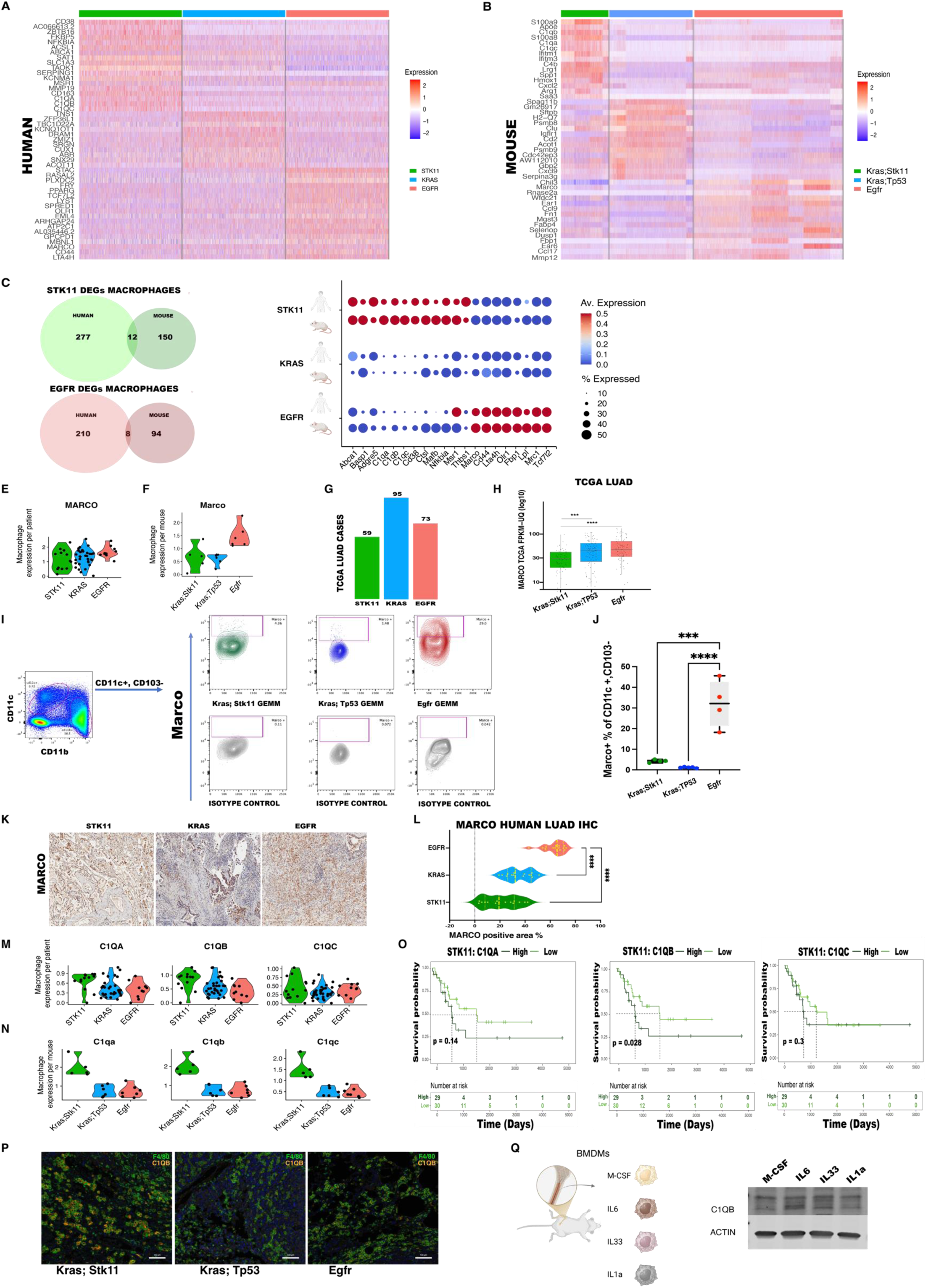
Genotype-guided profiling of LUAD macrophages reveals *MARCO* and *C1Q* as genotype-specific macrophage genes. **A**) Heatmap displaying macrophage DEGs among the three oncogenic subsets in human lung adenocarcinoma. **B**) Heatmap displaying macrophage DEGs among the three oncogenic subsets in mouse lung adenocarcinoma. **C**) Venn diagrams reveal shared human and mouse *STK11*-specific and *EGFR*-driven DEGs in lung adenocarcinoma macrophages. **D**) Dotplots verify *STK11*-specific expression of the twelve common cross-species genes and eight *EGFR*-specific genes in lung adenocarcinoma macrophages. **E**) Violin plots depicting average per-patient *MARCO* expression in human LUAD TAMs grouped by oncogenic driver mutations. **F**) Violin plots depicting average per-mouse *MARCO* expression in mouse LUAD TAMs stratified by genotype. **G**) TCGA LUAD cases stratification according to their genotype. **H**) Box plots of TCGA bulk RNA-seq *MARCO* expression comparison between the three major lung adenocarcinoma subtypes. **I**) Representative flow cytometry plots displaying MARCO expression in CD11C+ mouse myeloid populations in the three LUAD GEMMS. **J**) Box plots displaying the comparison of flow cytometry expression of *MARCO* as a percentage of CD11C+ myeloid cells among the three LUAD GEMMs. **K**) Representative images of *MARCO* IHC in patients with LUAD. Scale bars, 300 μm. **L**) MARCO IHC quantification of human LUAD patients. **M**) Violin plots comparing the average per-patient gene expression of *C1QA*, *C1QB*, and *C1QC* between human macrophages from LUAD patients with different oncogenic mutations. **N**) Violin plots comparing the average per-mouse expression of *C1qa*, *C1qb*, and *C1qc* between mouse macrophages of the three LUAD GEMMs subsets used in this study. **O**) Kaplan-Meier survival curves of patients with *STK11*-mutated lung adenocarcinoma using TCGA mRNA expression datasets for *C1QA*, *C1QB*, and *C1QB*. **P)** Multiplex immunofluorescence representative images validate the higher expression of C1QB in F4/80+ macrophages in *KRAS/STK11* mouse lung adenocarcinoma tumors. Scale bars, 100 μm. **Q**) Schematic depiction of mouse bone marrow-derived macrophage experiments and C1QB immunoblot indicating the IL6-dependent complement induction. Actin was used as a control for equal protein loading.

We then compared the human DEGs with those from the GEMMs and identified twelve *STK11*-associated and eight *EGFR-*associated common cross-species genes. This analysis did not yield any cross-species gene overlap in the *KRAS* group, most likely because of the presence of *KRAS* mutations in many of the *STK11* group human samples **(Figs. 4C and 4D)**. To verify that our observations were not driven by one specimen, we also assessed the average expression of *STK11* and *EGFR* TAM DEGs per specimen in our human and mouse TAM datasets **(Supplementary Fig. S6A)**.

Next, we sought to validate some of our *EGFR*-enriched macrophage genes, focusing on those with therapeutic potential. Macrophage receptor with collagenous structure (*MARCO*) has been proposed as a TAM immunotherapy biomarker in previous studies (40, 41). Our analysis suggests that *MARCO* could be a target, as it was consistently overexpressed in both *EGFR-*mutated patients and mice (**Figs. 4E and 4F**). To estimate the general pattern of *MARCO* expression across patients, we utilized the TCGA LUAD bulk RNA-seq data cohort grouped by oncogenic mutation status (**Fig. 4G and Supplementary Table S3**). Consistent with our earlier findings, *MARCO* expression was significantly higher in the TCGA *EGFR*-mutated group (**Fig. 4H**). We further validated the EGFR-associated enrichment of MARCO at the protein level by flow cytometry analysis of the macrophage compartment in the three GEMM models (**Fig. 4I**). Notably, Marco expression was markedly elevated in CD11c+ alveolar macrophages in the *EGFR*-mutated GEMM samples (**Fig. 4J**). In contrast, it was absent in CD11b+ myelomonocytic populations in all three GEMMs (**Supplementary Fig. S6B**), indicating that Marco expression was restricted to tissue-resident macrophages. Next, we sought to validate Marco using immunohistochemistry (IHC) in human LUAD samples with *STK11, KRAS,* and *EGFR* oncogenic mutations (**Fig. 4K**). We quantified five independent areas in samples from nine patients with LUAD (three patients from each LUAD genotype) and confirmed that Marco was enriched in *EGFR*-mutated human LUAD specimens (**Fig. 4L**).

Next, we focused on the twelve *STK11*-associated TAM genes to identify potential therapeutic targets, noting a significant enrichment of *C1Q* genes, such as *C1QA, C1QB,* and *C1QC*, which are key components of the classical complement pathway. Building on our earlier findings of phagocytosis enrichment and expression of the complement gene signature in *STK11*-mutated LUAD, we investigated the expression of *C1Q* genes in *STK11*-mutated LUAD TAMs. To confirm that the elevation of these genes in the *STK11*-mutated cohort represented a consistent trend rather than being driven by outlier samples, we analyzed the mean expression per patient in the human LUAD macrophage cohort (**Fig. 4M**) and the mean expression per mouse in the mouse LUAD macrophage cohort (**Fig. 4N**). Survival analysis of the TCGA LUAD *STK11*-mutated cohort revealed that high *C1QA*, *C1QB*, and *C1QC* gene expression was negatively associated with survival (**Fig. 4O**). To verify *C1Q* expression at the protein level, we performed multiplex immunofluorescence on lung tumor specimens from the three LUAD GEMMs, which verified the higher expression of C1QB^+^ among the F4/80^+^ macrophages in the *STK11*-mutated specimens (**Fig. 4P**).

Finally, to investigate the mechanism by which C1Q genes are upregulated in *STK11*-mutated TAMs, we isolated bone marrow-derived cells from 6- to 8-week-old mice. These cells were differentiated into macrophages over a five-day incubation period in the presence of macrophage colony-stimulating factor (M-CSF). Post-differentiation, macrophages were exposed to interleukin-6 (IL-6), interleukin-33 (IL-33), or interleukin-1 alpha (IL-1α) cytokines that were previously identified to be markedly enriched within *Stk11*-deficient tumors compared to *Kras* and *Kras; Tp53* tumor environments (13). Strikingly, a 72-hour IL-6 stimulation yielded a pronounced two-fold elevation in C1QB protein levels, underscoring IL-6’s putative role in orchestrating *C1QB* expression within the macrophage population (**Fig. 4Q and Supplementary Fig. S6C**).

In summary, our cross-species macrophage analyses identified *C1Q* and *MARCO* as distinguishing genetic features of *STK11-* and *EGFR*-mutated TAMs, underscoring their potential as genotype-specific immunotherapeutic targets for the treatment of LUAD.

### Oncogenic driver-specific transcriptomic analysis reveals cross-species conserved genes in LUAD epithelium

To gain insights into the genotype-driven gene signatures in LUAD epithelial cells and identify novel genotype-driven vulnerabilities, we stratified epithelial cells based on their oncogenic mutations in both human (**Fig. 5A and Supplementary Fig. S7A**), and mouse (**Fig. 5B**) samples. Most human epithelial cells were EpCAM+, with the majority identified as AT2 cells based on *SFTPB* expression. The smaller populations included AT1 (AGER+) and ciliated cells (FOXJ1+) (**Supplementary Fig. S7B**). All mouse epithelial cells were EpCAM+, and the majority were AT1/AT2, as verified by *Ager* and *Sftpb* expression, followed by a ciliated cluster (Foxj1+) (**Fig. S8A**). In our study, we analyzed the epithelial compartment, which consists of malignant and tumor-involved epithelial cells. We validated the presence of malignant cells in our samples by inference of copy number variation (Infer-CNV) analysis (**Supplementary Figs. S7C and S8B**) and histological analyses (**Supplementary Figs. S9A, S9B, S10A, and S10B**) of human and mouse samples.

**Figure 5.**
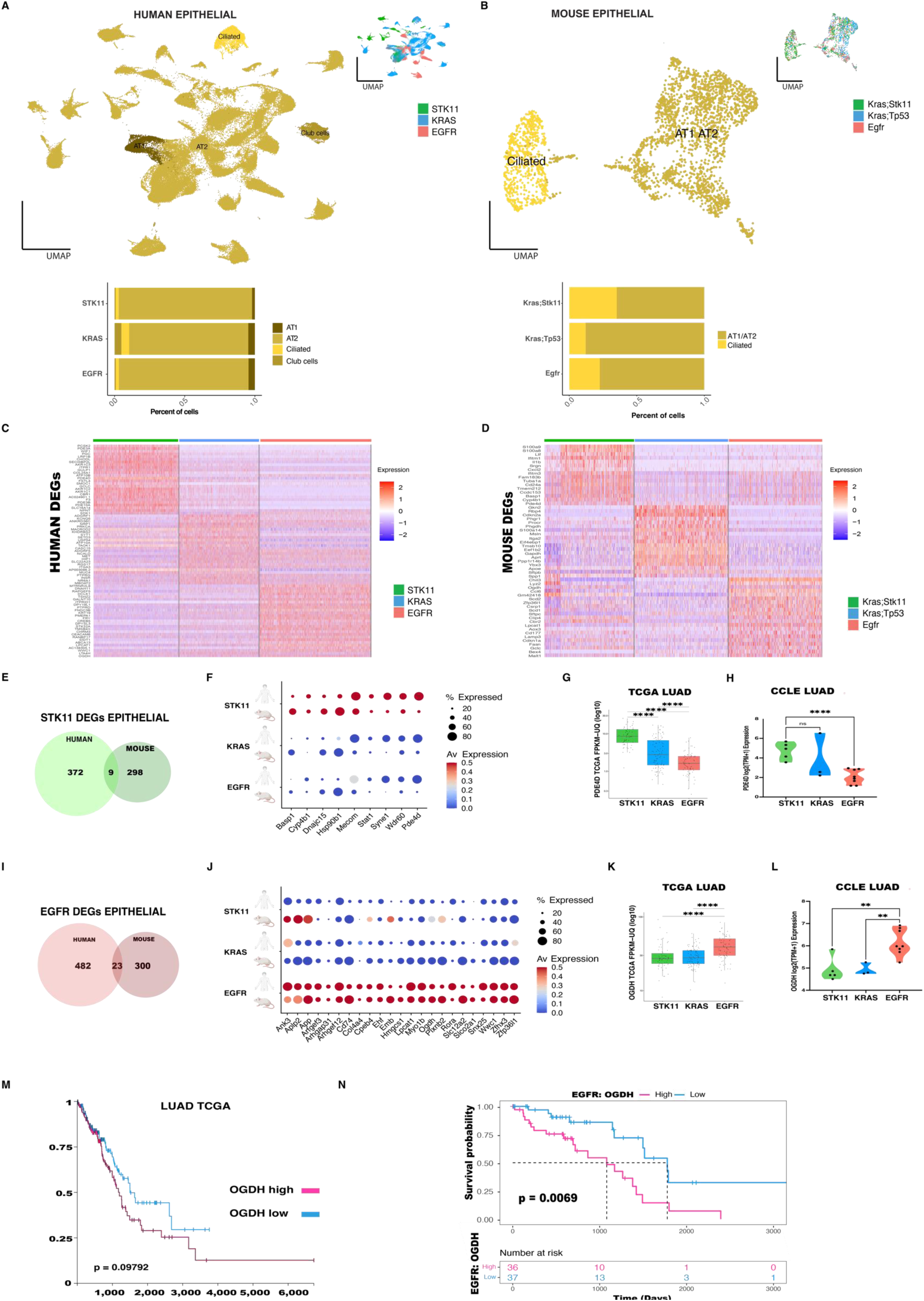
Oncogenic driver-specific transcriptomic analysis reveals conserved genes in LUAD epithelium. **A)** UMAP visualization and stacked plots of human epithelial cells from patients with LUAD grouped by three major oncogenic genotypes. **B)** UMAP visualization and stacked plots of mouse epithelial cells from LUAD GEMMs grouped by three key oncogenic genotypes. **C)** Heatmap of the top differentially expressed genes between epithelial cells from patients with LUAD harboring the three oncogenic mutations. **D)** Heatmap analysis showing the top differentially expressed genes between the epithelial cells of LUAD GEMMs harboring the three oncogenic mutations. **E)** Venn diagrams reveal common human and mouse *STK11*-specific genes in epithelial cells. **F)** Dotplots of the *STK11*-specific expression of the nine overlapping genes in epithelial cells. **G)** Box plot comparison of PDE4D mRNA expression among different LUAD genotypes from the TCGA dataset. **H)** Violin plots comparing PDE4D mRNA expression among different LUAD genotypes in human cell lines from the CCLE database. **I)** Venn diagrams reveal common human and mouse *EGFR*-specific differentially-expressed genes in epithelial cells. **J)** Bubble plots verify *EGFR*-specific expression of twenty-three overlapping genes in epithelial cells. **K)** Box plot comparison of OGDH mRNA expression among different LUAD genotypes from the TCGA dataset. **L)** Violin plots comparing OGDH mRNA expression among different LUAD genotypes in human cell lines from the CCLE database. **M)** Kaplan-Meier survival curve utilizing the TCGA lung adenocarcinoma mRNA expression dataset for the *OGDH* gene. **N)** Kaplan-Meier survival curve using the TCGA *EGFR*-mutated lung adenocarcinoma mRNA expression dataset for the *OGDH* gene.

Next, we sought to identify novel genotype-associated vulnerabilities in the epithelial compartment. We identified DEGs among the three LUAD genotypes in humans (**Fig. 5C an**d **Supplementary Table S4**) and mice (**Fig. 5D and Supplementary Table S5**).

Cross-examination of the human and mouse *STK11*-mutated epithelial DEGs yielded nine common genes (**Figs. 5E and 5F**), which included previously reported *STK11*-specific vulnerabilities, such as *HSP90B1* (19). Subsequently, we investigated the expression patterns of the nine genes in the TCGA LUAD cohort with available oncogenic mutations that we utilized previously. *PDE4D* expression was higher in the STK11 mutated group (**Fig. 5G**). This pattern was also verified in human LUAD cell lines, where *STK11*-mutated cell lines exhibited higher *PDE4D* expression than *KRAS-* and *EGFR*-mutated lines (**Fig. 5H**).

PDE4D was previously associated with several human cancers such as colon (29), prostate(30) and melanoma(31)

Moreover, we identified twenty-three common human-mouse DEGs markedly upregulated in the *EGFR*-mutated genotype (**Figs. 5I and 5J**). *CD74* (32) and *LPCAT1* expression have been previously associated with EGFR signaling in lung cancer (33) and glioblastoma (34). As described previously, we assessed the expression patterns of all identified genotype-specific genes in each patient (**Supplementary Fig. S8D**).

Extending our analysis, we investigated the expression of these twenty-three genes in the TCGA LUAD cohort. Stratifying TCGA LUAD patients by mutation status confirmed that *OGDH* expression was the highest in the *EGFR*-mutated group (**Fig. 5K**). Analysis of the cancer cell line encyclopedia (CCLE) dataset corroborated these findings, showing significantly higher *OGDH* expression in *EGFR*-mutated LUAD cell lines **(Fig. 5L**). Elevated OGDH expression was associated with poorer overall survival in LUAD patients from the TCGA cohort **(Fig. 5M),** with this association reaching greater statistical significance within the EGFR-mutant subset **(Fig. 5N).**

OGDH has been described as an actionable target in acute myeloid leukemia (35), glioblastoma (36) and gastric cancer(37).

Collectively, our comprehensive cross-species analyses of the epithelial compartment revealed PDE4D and OGDH as novel genotype-driven genes that are compelling candidates for targeted therapeutic intervention.

### Uncovering *PDE4D* and *OGDH* as genotype-specific therapeutic nodes in epithelial LUAD

Our cross-species analyses of epithelial cells unveiled *PDE4D* and *OGDH* as genotype-specific vulnerabilities, with expression selectively enriched in STK11-and EGFR-mutated LUAD, respectively. Importantly, elevated expression of these genes correlated with poorer clinical outcomes among LUAD patients, highlighting their clinical significance.

Building upon these findings, we systematically evaluated PDE4D expression across epithelial cells in LUAD patients and observed significantly higher expression levels within the STK11-mutated cohort compared to KRAS- and EGFR-mutant groups (**Fig. 6A**). To validate these observations at the protein level, we conducted immunohistochemistry (IHC) on LUAD specimens stratified by mutation status, confirming a pronounced enrichment of PDE4D in STK11-mutated tumors (**Fig. 6B and Supplementary Fig. S9A**). Concordantly, in genetically engineered mouse models (GEMMs) of LUAD, elevated PDE4D expression was similarly restricted to epithelial cells harboring STK11 mutations (**Fig. 6C**), a finding supported by IHC quantification (**Fig. 6D and Supplementary Fig. S9B**).

**Figure 6.**
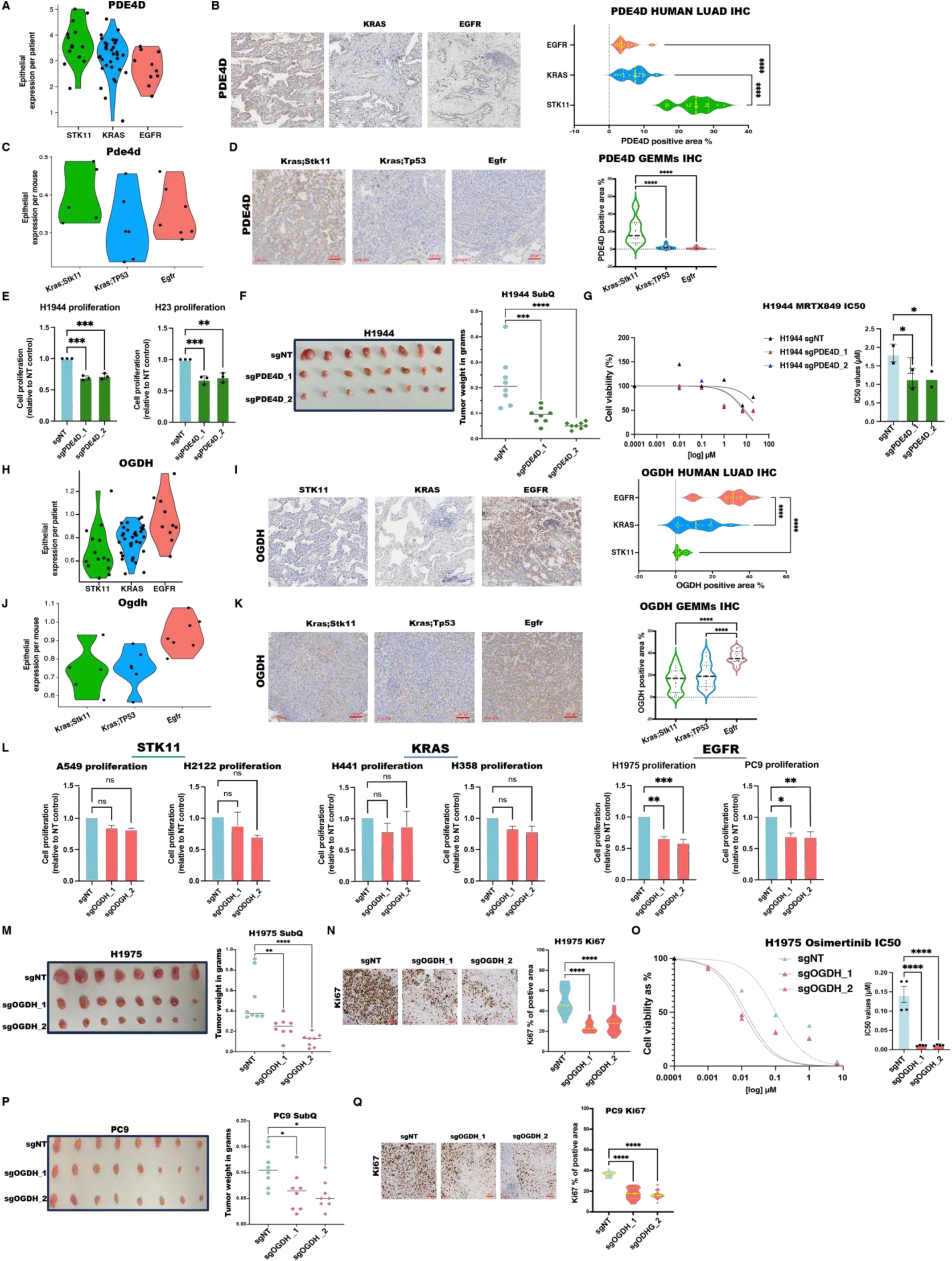
Uncovering *PDE4D* and *OGDH* as genotype-specific therapeutic nodes in epithelial LUAD. **A)** Comparison of average per-patient PDE4D mRNA expression among different human LUAD genotypes in scRNA-seq data from epithelial cells. **B)** Representative immunohistochemical staining of PDE4D in human LUAD specimens stratified by oncogenic genotype, accompanied by violin plots quantifying PDE4D expression across genotypes. Scale bars, 300 μm. **C)** Violin plots comparing the average per mouse PDE4D mRNA expression in the epithelial cells of the three LUAD GEMMs used in this study. **D)** Representative images of PDE4D IHC of the three LUAD GEMMs and violin plots comparing the PDE4D IHC quantification. Scale bars,100 μm. **E)** *In vitro* 2D assays comparing cell proliferation in H1944 and H23 LUAD cell lines upon CRISPR-mediated ablation of *PDE4D* (sgNT served as a non-targeting gRNA control). **F)** Representative images and corresponding quantitative analysis of tumor mass from subcutaneous H1944 xenografts following genetic ablation of *PDE4D.* **G)** Dose-response curves illustrate the sensitization of H1944 cells to MRTX849 upon *PDE4D* ablation seven days post-treatment and MRTX IC50s comparison upon *PDE4D* ablation. **H)** Comparison of the average per-patient OGDH mRNA expression among different human LUAD genotypes in the scRNA-seq data of epithelial cells. **I)** Representative immunohistochemical staining of OGDH in human LUAD specimens stratified by oncogenic genotype, accompanied by violin plots quantifying OGDH expression across genotypes. Scale bars, 300 μm. **J)** Violin plots comparing the average OGDH mRNA expression per mouse in epithelial cells of the three LUAD GEMMs used in this study. **K)** Representative images of OGDH IHC in LUAD GEMMs from different genetic backgrounds and violin plots comparing the OGDH IHC quantification. Scale bars, 100 μm. **L)** *In vitro* 2D assays comparing cell proliferation in various human LUAD cell lines (H1975, PC9, H441, H358, A549, H2212) upon *OGDH* CRISPR-mediated ablation (sgNT serves as a non-targeting gRNA control). **M)** Representative images and corresponding quantitative analysis of tumor mass from subcutaneous H1975 xenografts following genetic ablation of *OGDH*. **N)** Representative images of Ki67 IHC staining in H1975 tumors and violin plots comparing Ki67 staining in H1975 tumors. Scale bars are 100 μm. **O)** Dose-response curves illustrate the sensitization of H1975 cells to Osimertinib upon *OGDH* ablation 72 h post-treatment, after OGDH ablation and Osimertinib IC50s compared to *OGDH* ablation. **P)** Representative images and corresponding quantitative analysis of tumor mass from subcutaneous PC9 xenografts following genetic ablation of *OGDH*. **Q)** Representative images of Ki67 IHC staining in PC9 tumors and violin plots comparing Ki67 staining in PC9 tumors after *OGDH* ablation.

To dissect the relationship between PDE4D expression and STK11 mutational status, we next performed immunoblot analyses across a panel of human LUAD cell lines. Among KRAS/STK11-mutant lines, H1944 cells exhibited the highest PDE4D protein expression, followed by H23 and A549. In contrast, KRAS/TP53- and EGFR-mutant cell lines displayed minimal or undetectable PDE4D expression, except H358, which demonstrated only marginal levels (**Supplementary Fig. S9C**).

Functional interrogation of PDE4D dependency through CRISPR-mediated knockout revealed a profound impairment of proliferation in H1944 and H23 cells (**Fig. 6E and Supplementary Fig. S9D**). Extending these findings *in vivo*, PDE4D-deficient H1944 xenografts exhibited markedly suppressed tumor growth compared to controls (**Fig. 6F**). To further explore the translational potential of targeting *PDE4D*, we treated H1944 cells with MRTX849, an FDA-approved KRAS inhibitor. Notably, *PDE4D* loss significantly enhanced sensitivity to KRAS inhibition (**Fig. 6G**), suggesting a combinatorial therapeutic strategy for STK11-mutated LUAD.

Turning our attention to *OGDH*, analyses of epithelial cell expression profiles revealed preferential upregulation of *OGDH* in EGFR-mutated LUAD patients compared to KRAS- or STK11-mutated counterparts (**Fig. 6H**). This genotype-specific expression pattern was corroborated at the protein level via IHC in clinical LUAD specimens (**Fig. 6I and Supplementary Fig. S10A**), and was consistently recapitulated in GEMM models (**Figs. 6J**, **6K and Supplementary Fig. S10B**).

To assess the functional consequences of OGDH perturbation, we performed CRISPR-mediated knockout in EGFR-mutated LUAD cell lines. Ablation of OGDH significantly attenuated proliferation *in vitro* (**Fig. 6L and Supplementary Fig. S10C**). Consistent with these findings, subcutaneous xenografts of *OGDH*-deficient H1975 and PC9 cells displayed impaired tumor growth (**Figs. 6M and** 6**P)**, supporting an oncogenic role for OGDH in EGFR-driven LUAD. Recognizing the urgent need for improved therapies in EGFR-mutated LUAD, we next interrogated the therapeutic synergy between OGDH loss and standard-of-care tyrosine kinase inhibitors (TKIs). Remarkably, OGDH ablation sensitized H1975 cells to Osimertinib, significantly reducing the IC50 of the treatment (**Fig. 6O**). In further support, IHC analysis of Ki67 staining demonstrated a notable decrease in tumor cell proliferation following OGDH knockout in xenograft models (**Fig. 6N and Fig. 6Q)**

These findings unveil *PDE4D* and *OGDH* as compelling, clinically actionable vulnerabilities in STK11- and EGFR-mutated LUAD, respectively. Our results not only illuminate critical biological dependencies but also nominate these genes as promising therapeutic targets, underscoring the urgent need for translational efforts to exploit these novel susceptibilities to enhance therapeutic efficacy in molecularly defined LUAD subsets.

## Discussion

This study defines a comprehensive framework for dissecting the oncogenotype-specific ecosystems of LUAD, illustrating how driver mutations shape both tumor-intrinsic programs and the tumor microenvironment. Through integrative cross-species single-cell transcriptomics of 57 human LUAD specimens and 18 tumors from genetically engineered mouse models, we demonstrate that mutations in *STK11*, *KRAS*, and *EGFR* not only define malignant cell identity but also enact profound and lineage-specific rewiring of the tumor microenvironment, culminating in genotype-specific cellular compositions, transcriptional states, and therapeutic susceptibilities.

We propose a genetic seesaw model **(Fig. 7)** in which LUAD subtypes exist along a spectrum of immunologic states, with KRAS-mutant tumors occupying a transitional equilibrium, while STK11 and EGFR mutations define divergent immune microenvironments.

**Figure 7.**
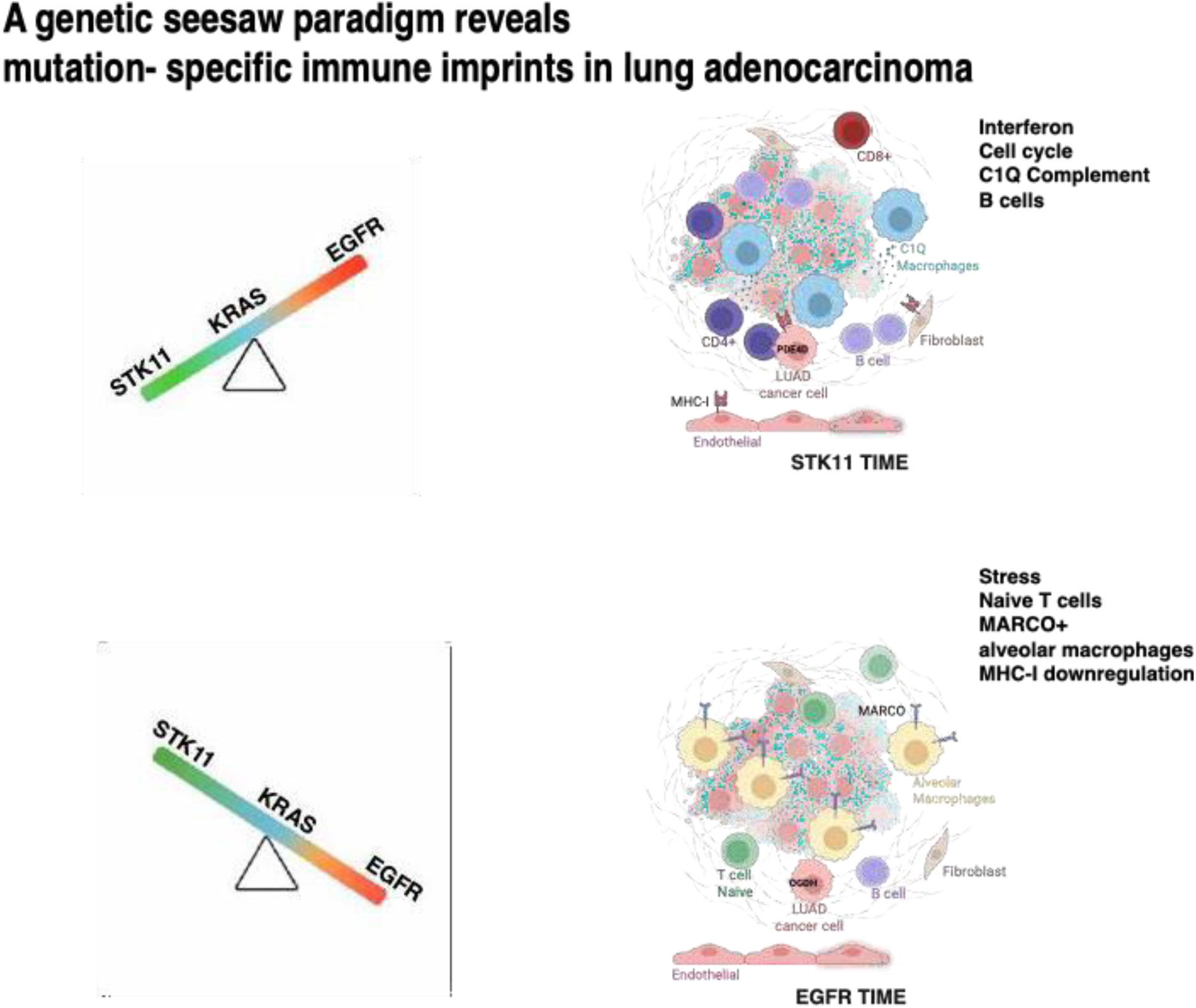

STK11-mutated tumors are characterized by an interferon- and complement-enriched, yet immunosuppressive, milieu. This is conserved across species and validated through transcriptomics, flow cytometry, and multiplex immunofluorescence. We identify C1Q complement components as key STK11-associated tumor-associated macrophage (TAM) genes, with high expression correlating with worse survival. The complement system, a core element of innate immunity, is increasingly recognized as a key player in the TME (38), (39). Recent studies emphasize that stromal-derived C1Q contributes to tumor progression and shapes tumor immunity in several human cancers, including adenocarcinomas of the colon, lung, breast, pancreas, and skin (40),(41), (42). Complement gene signatures are linked to tumor progression and poor prognosis in LUAD (43) ; however, their connection to STK11 loss has not been previously established. Mechanistically, we show that IL-6, abundantly secreted by STK11-deficient tumor epithelium, induces C1Q gene expression in macrophages, linking inflammatory cytokine signaling to complement activation. Given the increasing interest in complement-targeted therapies (44), (45), our data nominate the IL-6–C1Q axis as a promising target in STK11-driven LUAD, a subtype historically refractory to immunotherapy.

In contrast, EGFR-mutated tumors display a TME rich in alveolar macrophages and stress-naïve T cell programs, underpinned by global downregulation of MHC-I and loss of complement-regulatory interactions. MARCO, a scavenger receptor implicated in immune evasion, emerges as a conserved EGFR-specific TAM marker across human and mouse tumors, and is supported by flow cytometry, IHC, and TCGA analysis. MARCO-targeting strategies have shown preclinical efficacy in other cancers, including breast cancer (46), and NSCLC, (47) melanoma (48), and glioblastoma (49), and may hold therapeutic promise in EGFR-driven LUAD. These data underscore the plasticity of the myeloid compartment and the potential of genotype-informed myeloid reprogramming.

Notably, the contrasting immune microenvironments observed in STK11- and EGFR-mutant LUAD mirror fundamental biochemical opposition between these two signaling pathways. STK11, through activation of AMPK, dampens EGFR signaling via phosphorylation and promotes receptor internalization and degradation (50). Conversely, EGFR activation drives anabolic growth via PI3K-AKT-mTOR signaling, opposing the energy-conserving, catabolic programs of the LKB1–AMPK axis. Consistent with this, EGFR and STK11 mutations are infrequently co-occurring in LUAD and may be mutually exclusive. In rare cases where they co-occur, STK11 loss has been associated with resistance to EGFR-targeted therapies (51), (52). These opposing roles are further reflected in the immune landscape: whereas STK11 mutations drive inflammatory signaling and complement activation, EGFR-mutant tumors exhibit profound immune suppression, with reduced antigen presentation and enrichment of macrophages expressing immune-dampening receptors like MARCO.

Importantly, our study also identifies metabolic and signaling vulnerabilities in the epithelial compartment that are aligned with oncogenic genotype. PDE4D, enriched in STK11-mutant LUAD, and OGDH, selectively upregulated in EGFR-mutant tumors, were validated across human and mouse tumors and functionally confirmed through CRISPR loss-of-function assays and xenografts. Both genes are associated with poor prognosis and sensitize LUAD cells to KRAS or EGFR inhibitors, respectively. These findings open new therapeutic avenues by coupling targeted therapy with metabolic inhibition in genotype-defined LUAD subtypes.

OGDH, a key enzyme of the tricarboxylic acid (TCA) cycle, has dual roles in energy metabolism and redox regulation(34). Our study reveals that EGFR-driven lung adenocarcinomas, OGDH overexpression within the epithelial compartment, coincides with the activation of stress-associated transcriptional programs, suggesting that its upregulation not only reflects but also reinforces a broader metabolic adaptation to an oxidative stress–enriched tumor immune microenvironment. Conversely, PDE4D regulates cAMP signaling, with known roles in modulating interferon and cytokine responses(53), (54). In the context of STK11-mutant LUAD, targeting PDE4D is maybe a tool to therapeutically recalibrate the hyperinflammatory and proliferative immune landscape.

Taken together, our work provides a foundational resource and conceptual advance for the field of cancer immunogenomics. It illustrates how oncogenic mutations propagate their effects beyond the confines of tumor cells to orchestrate an immunologic and stromal response that is both distinct and targetable. In the future, a more granular characterization of GEMMs across a spectrum of malignancies, coupled with cross-comparisons with publicly accessible human cancer scRNA atlases, can improve the identification of novel therapeutic targets. Moreover, this refined approach will diminish reliance on animal models for preclinical testing by focusing on more robust and biologically relevant targets. Finally, our study advocates for the routine incorporation of oncogenic genotype into TME-based therapeutic stratification and calls for the development of precision immunotherapies tailored to the mutational context of the tumor.

## Data Code and Availability

The sequencing data were deposited in NCBI: Human snRNA-seq (GEO: GSE279384) and mouse scRNA-seq (GEO: GSE279746).

## Acknowledgments

NYU Langone Preclinical Imaging Laboratory (partially funded by [P30CA016087](https://www.sciencedirect.com/science/article/pii/S1535610822005621?via%3Dihub&gs1) and [P41EB017183](https://www.sciencedirect.com/science/article/pii/S1535610822005621?via%3Dihub&gs1)) for MRI, Genome Technology Center (partially supported by [P30CA016087](https://www.sciencedirect.com/science/article/pii/S1535610822005621?via%3Dihub&gs2)) for library preparation and sequencing, and Experimental Pathology Research Laboratory (partially funded by [P30CA016087](https://www.sciencedirect.com/science/article/pii/S1535610822005621?via%3Dihub&gs3)) for immunohistochemistry. The NYULH Center for Biospecimen Research and Development, Histology, and Immunohistochemistry Laboratory (RRID: SCR_018304) is partially supported by the Laura and Isaac Perlmutter Cancer Center Support Grant (NIH/NCI P30CA016087).

Main funding source for this study: National Cancer Institute: R01CA293718

## Author Contributions

A.K., M.P., I.Y., and K.-K.W. supervised the work, designed all experiments, and interpreted the data. All the authors contributed to the experiments and manuscript preparation. A.K. and M.P. wrote I.Y., K.-K.W. edited the manuscript, and all authors approved the manuscript.

## Methods

### Animal studies

The Institutional Animal Care and Use Committee at the New York University Grossman School of Medicine reviewed and approved all animal studies. *EGFR*-mutant GEMMs harbor the conditional activating mutation L858R of the human *EGFR* gene incorporated into the *Col1a1* locus. In this model, Cre recombinase is induced by intranasal inhalation of 5 × 10^7^ plaque-forming units of adeno-Cre (University of Iowa adenoviral core), and tumor nodules typically appear 20– 25 weeks after induction.

*Kras* (*Kras*^LSL-G12D/+^) was crossed with *Stk11* or *Tp53* conditional knockout (*Stk11* fl/fl or *Tp53* fl/fl). In these models, Cre-recombinase is induced by intranasal inhalation of 1 × 10^7^ plaque-forming units of adeno-Cre (University of Iowa adenoviral core), and tumors typically appear 6 weeks after induction for the *Kras; Stk11* model and 8 weeks for the *Kras; Tp53* model.

### Generation of xenograft models

To generate subcutaneous xenografts, each mouse was inoculated with tumor cells in both flanks: 2 × 10^6^ for H1975 and PC9, and 4 × 10^6^ for H1944 in 200 μL of phosphate-buffered saline (PBS) per flank. Tumors were weighed at the endpoint.

### Human lung adenocarcinoma patients

#### Specimen collection

Snap-frozen stage I lung cancer and matching adjacent lung specimens (within the same lobe, segment, or wedge resection) from 56 patients undergoing R0 resection with lymph node dissection were prospectively collected and archived at -80 °C between 2005 and 2015 under an IRB-approved NYULH protocol (R8896). Subjects who received antibiotics (except perioperative), steroids, radiation, immunotherapy, or chemotherapy within one month before surgery were excluded from the study. Subjects were assessed postoperatively via an in-person clinic visit and surveillance chest CT every three months after surgery for two years, every six months for the third year, and annually thereafter.

#### SnRNA sequencing

Nuclei were prepared for 10x Genomics-based single-nuclei RNA-seq analysis. Briefly, each frozen sample was thawed and macerated in CST buffer for 10 minutes, filtered with a 70-μm pluriStrainer (pluriSelect, Cat #43-10070-40), and centrifuged at 500 × g for 5 min at 4 °C to pellet the nuclei. Nuclei were resuspended in the same buffer without detergent, filtered with a 10-μm pluriStrainer (pluriSelect, Cat #43-10070-40), and counted using AOPI on a Nexcelom Cellometer (Nexcelom Bioscience, Lawrence, MA). Approximately 10,000 nuclei were loaded immediately into each channel of a 10X Genomics Chromium chip, using 5-prime v1.1 chemistry according to the manufacturer’s protocol (10x Genomics, Cat #PN-1000165). The resulting cDNA and indexed libraries were checked for quality on an Agilent 4200 TapeStation (Agilent Technologies, Santa Clara, CA), and then quantified and pooled for sequencing on an Illumina NovaSeq 6000 (Illumina, San Diego, CA).

#### snRNA-seq analysis pipeline

Sequencing reads were trimmed with cutadapt and aligned to the GRCh38 reference transcriptome, including introns (ref data cellranger-GRCh38-3.0.0), using STARsolo v2.7.3a (55) .We identified nuclei-containing droplets using in-house tools that analyze knee plots (that depict the number of genes/transcripts detected in each nucleus vs. each nucleus’s ranking) for an inflection point and used a graph-based method to recover a portion of barcodes below the inflection point if their QC metrics are good. Ambient RNA contamination was removed from each sample’s count matrix using an in-house tool similar to FastCar (56). We further filtered nuclei assigned to each cell type compartment using compartment-specific thresholds for several detected genes. Doublets were identified and removed using an in-house implementation of the Solo algorithm (57). Batch correction and expression normalization were performed using scVI scvi-tools (58). Normalized expression values are represented as log10(Transcript Per 10 K). Finally, clustering was performed using the Leiden algorithm. For visualization purposes, we generated four low-dimensional embeddings: tSNE projection of all nuclei, UMAP of all nuclei, composite tSNE (merging individual tSNEs generated per compartment), and composite UMAP.

#### Immunohistochemistry

Rabbit polyclonal anti-human unconjugated oxoglutarate dehydrogenase (OGDH)(Atlas Antibodies, Cat #HPA020347), rabbit polyclonal anti-human unconjugated cAMP-specific 3’,5’-cyclic phosphodiesterase 4D (PDE4D) (Abcam, Cat #ab14613), and rabbit polyclonal anti-human unconjugated macrophage receptor with collagenous structure (MARCO) (Atlas Antibodies, Cat #HPA063793) were optimized using a Ventana Medical Systems Discovery Ultra platform (Ventana Medical Systems, Oro Valley, AZ). Briefly, formalin-fixed, paraffin-embedded, 5-µm human archival tissue sections were prepared on SlideMate plus microscope slides (Cat #TT-4041I-PS-8218-1) and stored at room temperature before use. The sections were deparaffinized online using Discovery Wash (Ventana Medical Systems, Cat #950-510) for 24 min at 69 °C. Endogenous peroxidase was blocked using 3% hydrogen peroxide for 4 min. Sections were antigen-retrieved using Ultra Cell Conditioner 1 (Ventana Medical Systems, Cat #950-224) as follows: OGDH was retrieved for 24 min at 100 °C, PDE4D was retrieved for 92 min at 100 °C, and MARCO was retrieved for 24 min at 100 °C. Each antibody was diluted (Ventana Medical Systems, Cat #ADB250) and incubated as follows: OGDH, 1:400, incubated for 6 h; PDE4D, 1:50, incubated for 12 h; and MARCO, 1:100, incubated for 3 h. Antibodies were detected using a goat anti-rabbit HRP-conjugated multimer and 3,3-diaminobenzidine (DAB) chromogen. The slides were washed in distilled water, counterstained with hematoxylin, dehydrated through graded alcohols, cleared in xylene, and mounted with synthetic permanent medium. Appropriate positive and negative controls were used in this study.

### Mouse lung single-cell preparation

The mice were euthanized, and the lungs were perfused with sterile PBS from the left ventricle. Whole lungs were minced and digested with collagenase D (Roche, Cat #11088866001) and DNase I (Roche, Cat #10104159001) in Hank’s balanced salt solution at 37 °C for 30 min. After incubation, the digested tissue was filtered through a 70-μm cell strainer (Thermo Fisher Scientific, Waltham, MA, USA) to obtain single-cell suspensions. Single cells were treated with Red Blood Cell lysis buffer (BioLegend, Cat #420301) to lyse red blood cells for 5 minutes at room temperature. Subsequently, the samples were stained with hashing antibodies (BioLegend, Totalseq B anti-mouse Hashtags) for 30 min on ice. DAPI (BioLegend; Cat #422801) was used for live/dead separation at a final concentration of 1 ng/mL.

### Single-cell RNA sequencing

Single-cell suspensions were prepared as described above and sorted using live/dead DAPI staining. The cells were then resuspended into single cells at 1 × 10^6^ cells/mL in PBS with 0.4% BSA for 10x Genomics processing. Cell suspensions were loaded onto a 10x Genomics Chromium instrument to generate single-cell gel beads in an emulsion (GEM). Approximately 30,000–40,000 cells are loaded per channel. scRNA-seq libraries were prepared using the following: Single Cell 3’ Reagent Kits: Chromium Single Cell 3’ Library and Gel Bead Kit v2 (10x Genomics, Cat #PN-120237), Single Cell 3’ Chip Kit v2 (10x Genomics, Cat #PN-120236), and i7 Multiplex Kit (10x Genomics, Cat #PN-120262) according to the manufacturer’s protocol, followed by the Single Cell 3’ Reagent Kits v2 User Guide (Manual Part #CG00052 Rev A). Libraries were sequenced on an Illumina NovaSeq 6000 with 2 × 150 paired-end reads for >90% sequencing saturation.

### Single-cell sequencing analysis

Cells with low-quality transcriptomes were excluded from the mouse scRNA-seq data analysis. Specifically, cells with fewer than 700 or more than 6,000 UMIs or more than 20% mitochondrial or ribosomal reads were filtered out. The data were processed for single-cell transformation using Seurat v4.0.3 (59).

The Seurat function FindAllMarkers was used for DEG comparisons between the genotypes. We presented the top 15–20 average_log2FC genes, whereas we used all significant DEGs (varying between 50 and 250) for analysis.

For the GSE analysis, all DEGs were converted to humans using the useMart function and compared with the hallmark dataset using the RITAN R package.

For gene module analysis, we scored all human cells from each cell type to the list of appropriate gene programs from Gavish et al, using the Seurat AddModuleScore function. Local mutation mixing entropy analysis was calculated for each cell type within the TIME compartment of the mouse dataset, computing Shannon entropy on the k-nearest neighbors in UMAP space, using the FNN and the entropy functions.

### Cell type identification

The Seurat package was used to cluster the cells with a resolution of 0.2 and to identify differentially expressed genes between the resulting clusters based on the average log2 fold change (FC). These genes were then cross-referenced with existing literature to identify markers for various immune and non-immune cell types, including PTPRC (immune cells), CD3E (T cells), CD19 (B cells), CSF1R (macrophages/monocytes), NKG7 (NK cells), PECAM1 (endothelial cells), COL1A1 (fibroblasts), EPCAM, and KRT (epithelial cells).

### Mouse single-cell data integration

To correct for the batch effect of three separate experiments with multiple mice for each model in each experiment, we first integrated the gene expression matrices for each genotype across experiments together, using the ‘IntegrateData’ function from Seurat, and the top differentially expressed 8,000 genes. We used this integrated data from each genotype to annotate the cell types based on marker expression and clustering. To allow cross-species comparison and joint presentation of the data across the genotypes, we then integrated the raw counts data from the three genotypes across experiments, again using the Seurat ‘Integrate Data’ function. We projected the cell type annotation from the first approach separately on each model, which resulted in similar cell type clusters when visualizing the joint integration data. All downstream analyses and comparisons of specific genes or modules across genotypes were performed on the unintegrated data.

### Cell-type averaging RNA-seq analysis

To study per-patient or per-mouse differences across genotypes, we averaged the gene expression data of every annotated cell type separately for each patient or mouse as step one. For Figure 1 global analysis, we averaged all the cell type-specific pseudo-bulks for each patient or mouse from step one, creating one value for each patient or mouse while correcting for differences in cell type composition between the individuals. For module score heatmaps, we averaged the relevant cell type-specific pseudo-bulks for each patient or mouse from step one across all individual mice or patients of the same genotype, creating one value for each genotype from each species while correcting for differences in cell type abundance between the individuals.

### Cell-cell communication analyses

Intercellular communication analysis was performed using CellChat (v2.1.2), a computational framework designed to infer and visualize cell–cell communication networks from single-cell RNA sequencing (scRNA-seq) data. Normalized expression matrices and cell identities were exported from Seurat objects by following CellChat’s instructions. The computeCommunProb function was used to compute the communication probability and infer the cellular communication network. ‘TriMean’ was used for computing the average gene expression per cell group. Cell-cell communications with less than 10 cells in each cell group were filtered out from further consideration. Communication probabilities were aggregated at the signaling pathway level via aggregateNet. Network centrality and interaction strength were analyzed to infer dominant incoming and outgoing signaling roles for each cell type. All visualizations, including network plots and hierarchical signaling diagrams, were generated using CellChat’s native plotting functions and a customized R script.

### Transcription factor (TF) activity analysis

Gene regulatory network inference and transcription factor (TF) activity analysis were performed using pySCENIC (v0.12.1), a computational framework for identifying cell–state–specific regulatory programs from scRNA-seq data. Raw gene expression matrices were first normalized and filtered to retain genes with sufficient variability across cells. SCENIC was run using the default three-step pipeline: (1) co-expression modules were identified using the GRNBoost2 algorithm; (2) cis-regulatory motifs were detected within the co-expression modules using RcisTarget to refine regulons; and (3) regulon activity was quantified across single cells via the AUCell algorithm. This approach enabled the identification of active transcriptional programs and regulatory network dynamics across distinct cell populations. Heatmap visualization of cell types, regulon activity, and clustering was conducted using ggplot functions in R based on AUCell scores.

### Histology and immunohistochemistry for mouse specimens

For histological and immunohistochemical analyses, the lungs were initially perfused with 10% formalin, fixed for 48 h in formalin at 4 °C, washed with 70% ethanol, and then embedded in paraffin. Subsequently, 5-µm tissue sections were prepared from formalin-fixed paraffin-embedded tissues, which underwent immunoperoxidase analysis following a sequence of preparatory steps: baking at 60 °C for 1 h, deparaffinization, and rehydration through multiple immersions in 100% xylene (4 times, 3 min each) and 100% ethanol (4 times, 3 min each), followed by rinsing in running water for 5 min. To block peroxidase activity, the sections were treated with 3% hydrogen peroxide in methanol for 10 min and rinsed under running water for another 5 min. Antigen retrieval was performed using a pressure cooker and Target Retrieval Solution Citrate, pH 6.1 (Agilent Technologies; Cat #S1699) at 120 °C. After retrieval, the slides were cooled for 15 min before being transferred to Tris-buffered saline (TBS). The sections were incubated with primary antibodies (OGDH, PDE4D, and Ki-67) for 40 min at room temperature. The secondary antibody, Leica Novolink Polymer (Leica Biosystems, Cat #RE7161), was applied for 30 min. Incubations were carried out in a humidified chamber at room temperature, with rinses in TBS between each step. For visualization, sections were developed using 3,3’-diaminobenzidine (DAB) as the substrate and counterstained with Mayer’s hematoxylin. Immunohistochemistry images were analyzed and quantified using the FIJI software. Five independent areas from each (blinded) specimen were analyzed and quantified for the IHC comparative analyses.

### Isolation and differentiation of bone marrow-derived macrophages

Tibias and femurs were harvested from euthanized 6–8-week-old mice and excess soft tissue was removed. Both ends of each bone were cut, and the bone marrow (BM) was flushed out using 5 mL RPMI-1640 medium supplemented with 10% fetal bovine serum (FBS) and delivered through a 23-G needle. The cell suspension was filtered through a 70-μm cell strainer and centrifuged at 300 × g for 5 min at room temperature. The cell pellet was resuspended in 1 mL of ACK lysis buffer and incubated for 1 min to lyse the red blood cells. The cells were centrifuged at 300 × g for 5 min and resuspended at the desired concentration in RPMI-1640 medium for subsequent culture. Subsequently, a total of 10^7^ bone marrow-derived cells were resuspended in 10 mL RPMI-1640 medium containing 10% FBS, 1% L-glutamine, Primocin (Invitrogen, Cat #ant-pr-1), and 20 ng/mL murine recombinant M-CSF (Thermo Fisher Scientific, PeproTech, Cat #315-02-10UG). Cells were plated in tissue culture-treated petri dishes and incubated at 37 °C in a humidified 5% CO₂ atmosphere. On Day 3, the medium was replaced with fresh RPMI-1640 containing M-CSF. On Day 5, macrophages were stimulated for 72 h under one of the following conditions: (1) Control: M-CSF only; (2) IL-6: M-CSF + 100 ng/mL IL-6 (Peprotech, Cat #216-16); (3) IL-33: M-CSF + 100 ng/mL IL-33 (Peprotech, Cat #210-33); or (4) IL-1α: M-CSF + 100 ng/mL IL-1α (Peprotech, Cat #211-11A)After 72 h, macrophages from each treatment group were collected and prepared for immunoblot analysis to assess protein expression.

### Murine lung cell isolation for immune analysis

Mice were euthanized, and the lungs were perfused with sterile phosphate-buffered saline (PBS) via cardiac perfusion through the left ventricle to clear blood from the pulmonary vasculature. Whole lungs were excised, finely minced, and enzymatically digested in Hank’s Balanced Salt Solution (HBSS) containing collagenase D (Roche, Cat #11088866001) and DNase I (Roche, Cat #10104159001) at 37 °C for 30 min. After digestion, the lung tissue suspensions were filtered through a 70-μm cell strainer to obtain single-cell suspensions. The resulting cell suspension was treated with 1× Red Blood Lysis buffer (BioLegend, Cat #420301) to remove the residual red blood cells. Cell pellets were resuspended in PBS containing 2% FBS and stained with the indicated cell surface antibodies. The cells were fixed and permeabilized using a BD Cytofix/Cytoperm Fixation/Permeabilization Kit (BD Biosciences, Cat #554714). Stained cells were analyzed on a BD Biosciences LSR Fortessa flow cytometer, and data were processed using FlowJo software.

### Multiplex immunofluorescence

Paraffin blocks of tumor-bearing lungs were prepared as described above. Five-µm-thick sections were sliced from paraffin blocks and processed for staining. Sections were stained with hematoxylin and eosin or reagents from the Akoya Biosciences Opal multiplex automation kit (Akoya Biosciences, Cat #ARD1001EA) on a Leica BondRX, according to the manufacturer’s instructions.

The process included sequential epitope retrieval that was performed using Leica Biosystems solution ER2 (EDTA-based, pH 9, Cat # AR9640). This was followed by primary and secondary antibody incubation with Opal fluorophores (Op570 Akoya, Cat # FP1488001KT or Op690 Akoya, Cat #FP1497001KT) and tyramide signal amplification. Primary antibodies against mouse F4/80 (CST, Cat #70076, Clone D259R) and mouse C1QB (Thermo Fisher Scientific Technology, Cat # PA5-81321) were used.

Horseradish peroxidase-coupled secondary antibodies (Rabbit-on-Rodent HRP polymer; Biocare, Cat # RMR622) were used. After sequential heat retrieval, only the opal fluorophores remained covalently bonded to the epitope.

Sections were counterstained with DAPI (Cat #FP1490; Akoya Biosciences) and mounted using the Prolong Gold Antifade Reagent (Invitrogen, Cat #P36930). Semi-automated image acquisition was performed using an Akoya Vectra Polaris (PhenoImagerHT) multispectral imaging system at 20X magnification. Selected tissue fields were outlined manually for spectral unmixing using InForm version 2.4.11 software (Akoya Biosciences).

### 2D cell proliferation assays

One thousand cells were seeded in tissue culture-treated 96-well/12-well plates and incubated for three or seven days at 37 °C, depending on the cell line. If the media contained various concentrations of drugs, they were replaced with fresh drugs every three days. Cell viability was determined using a Cell Counting Kit-8 (Cat #ALX-850-039-0100; Enzo Life Sciences), according to the manufacturer’s protocol. Luminescence was measured at 450 nm for 4 h after adding CCK-8 reagent to the cells using a FlexStation 3 multi-mode microplate reader, according to the manufacturer’s instructions.

### Cell lines

All human LUAD cell lines were maintained in RPMI 1640 medium (Cat #11875119; Thermo Fisher Scientific) supplemented with 10% fetal bovine serum (FBS, Sigma-Aldrich, Cat #12306C), GlutaMAX Supplement (Thermo Fisher Scientific, Cat #35050061), and 1× antibiotic-antimycotic (Thermo Fisher Scientific, Cat #15240062). HEK-293T cells (ATCC, Cat #CRL-1573) were maintained in Dulbecco’s modified Eagle’s medium (DMEM, Thermo Fisher Scientific, Cat #11965118) supplemented with 10% FBS and 1× antibiotic-antimycotic solution. All cell lines were cultured at 37 °C in a humidified incubator containing 5% CO₂ and confirmed to be mycoplasma-free using the Universal Mycoplasma Detection Kit (ATCC, Cat #30-1012K).

### Cell line CRISPR knock-out generation

For lentivirus production, HEK-293T cells were co-transfected with Cas9 lentiCRISPR v2 puro (Addgene, Cat #52961) along with the packaging plasmids psPAX2 (Addgene, Cat #12260) and pMD2.G (Addgene, Cat #12259) using Lipofectamine 3000 (Invitrogen, Cat #L3000008), following the manufacturer’s protocol. Cell culture supernatants containing viral particles were collected 72 h later and purified through 0.45-μm filters (Corning, Cat #431225) to eliminate cellular debris. Subsequently, viruses were used to infect various cell lines in the following 1-hour spin-infection at 600 × g at 32 °C in the presence of 8 μg/mL polybrene (Sigma-Aldrich, Cat #TR-1003) and subsequent puromycin (Invivogen, Cat #ant-pr-1) selection 24 h post-infection.

### Immunoblotting

Cell pellets were lysed using RIPA buffer (Cat #89900, Thermo Fisher Scientific) with protease and phosphatase inhibitor cocktail (Cat #78440; Thermo Fisher Scientific). Proteins were quantified using a Pierce BCA Protein Assay Kit (Thermo Fisher Scientific, Cat #23225). Subsequently, equal quantities of total protein from each sample were mixed with 6x reducing Laemmli SDS sample buffer (Boston BioProducts Inc., Cat #BP-111RR) and distilled water and boiled at 95 °C for five minutes to achieve protein denaturation. Subsequently, the samples were loaded and electrophoresed on 4–20% Criterion TGX Precast Midi Protein gels (Bio-Rad, Cat #5671094) and then transferred onto nitrocellulose membranes for immunoblotting. The membranes were incubated overnight at 4 °C with primary antibodies targeting OGDH (Atlas Antibodies, Cat #HPA020347), PDE4D (Abcam; Cat #ab14613), C1QB (Sigma-Aldrich Cat #SAB1401034) and β-actin (Sigma-Aldrich, Cat #A5441). For detection, IRDye 800CW donkey anti-rabbit IgG (LI-COR Biosciences, Cat #926-32213) and IRDye 680RD donkey anti-mouse IgG (LI-COR Biosciences; Cat #926-68072) secondary antibodies were incubated for 1 h at room temperature, washed three times with PBS containing Tween 0.01%, and visualized using an Odyssey detection system (LI-COR Biosciences).

### Statistical analyses

GraphPad Prism 10 (GraphPad Software, La Jolla, CA) was used for all statistical analyses. Data were analyzed by Student’s t-test (two-tailed) and Analysis of Variance (ANOVA) for three or more groups. Error bars represent the standard error of the mean (SEM). *P < 0.05; **P < 0.01; ***P < 0.001; ****P < 0.0001.

### Illustration tool

All the schematic illustrations were created with BioRender (https://biorender.com/

---

**Figure S1.**
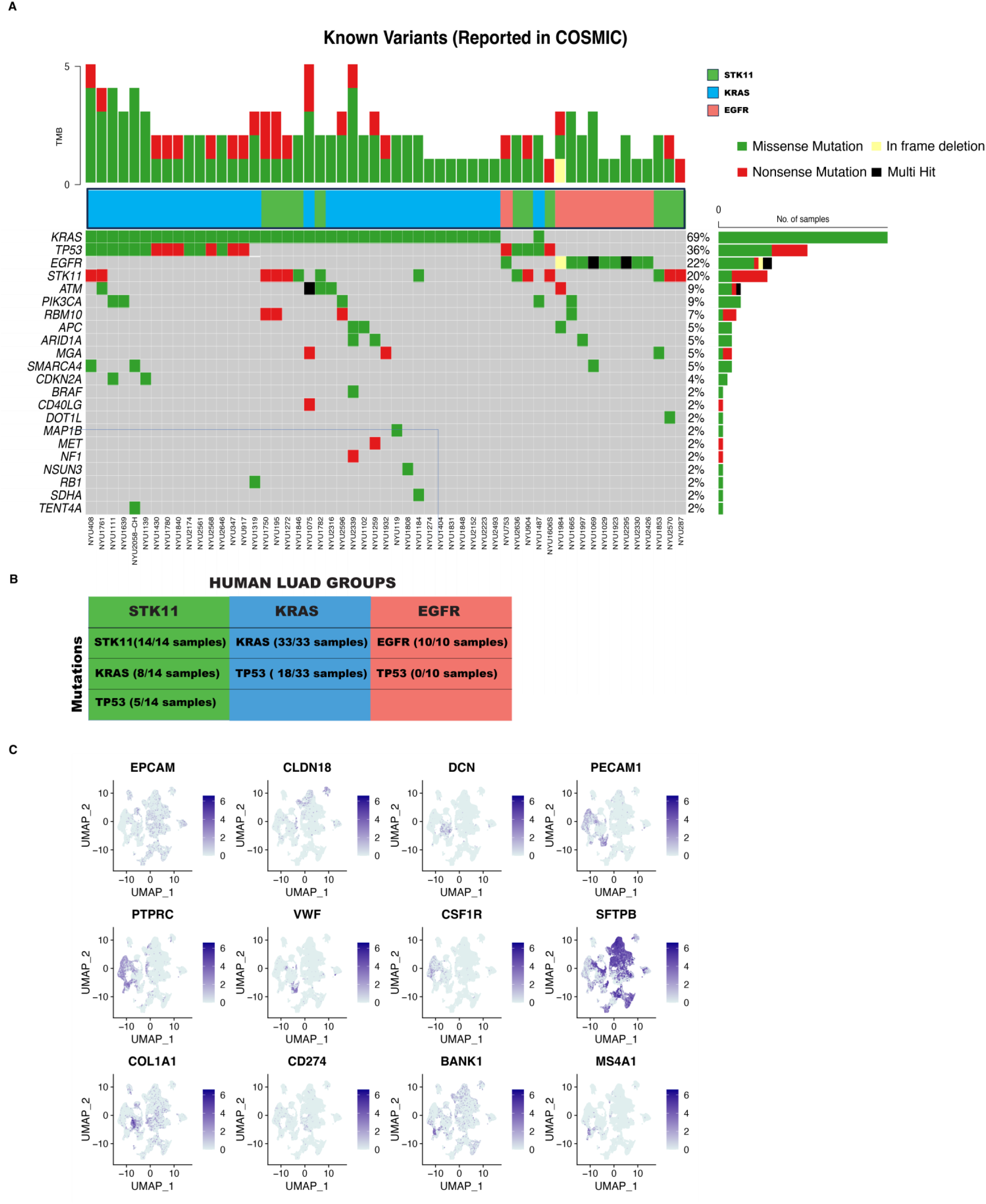
Single-nucleus RNA sequencing and DNA sequencing of frozen human lung adenocarcinoma specimens allows genotype-driven profiling. **A)** OncoPrint diagram of the mutation profiles of LUAD specimens used for snRNA-seq. **B)** Table depicting the Grouping of human LUAD samples based on the presence of major oncogenic mutations. **C)** UMAP visualization of general cell-type-specific human gene markers used to identify the cell types.

**Figure S2.**
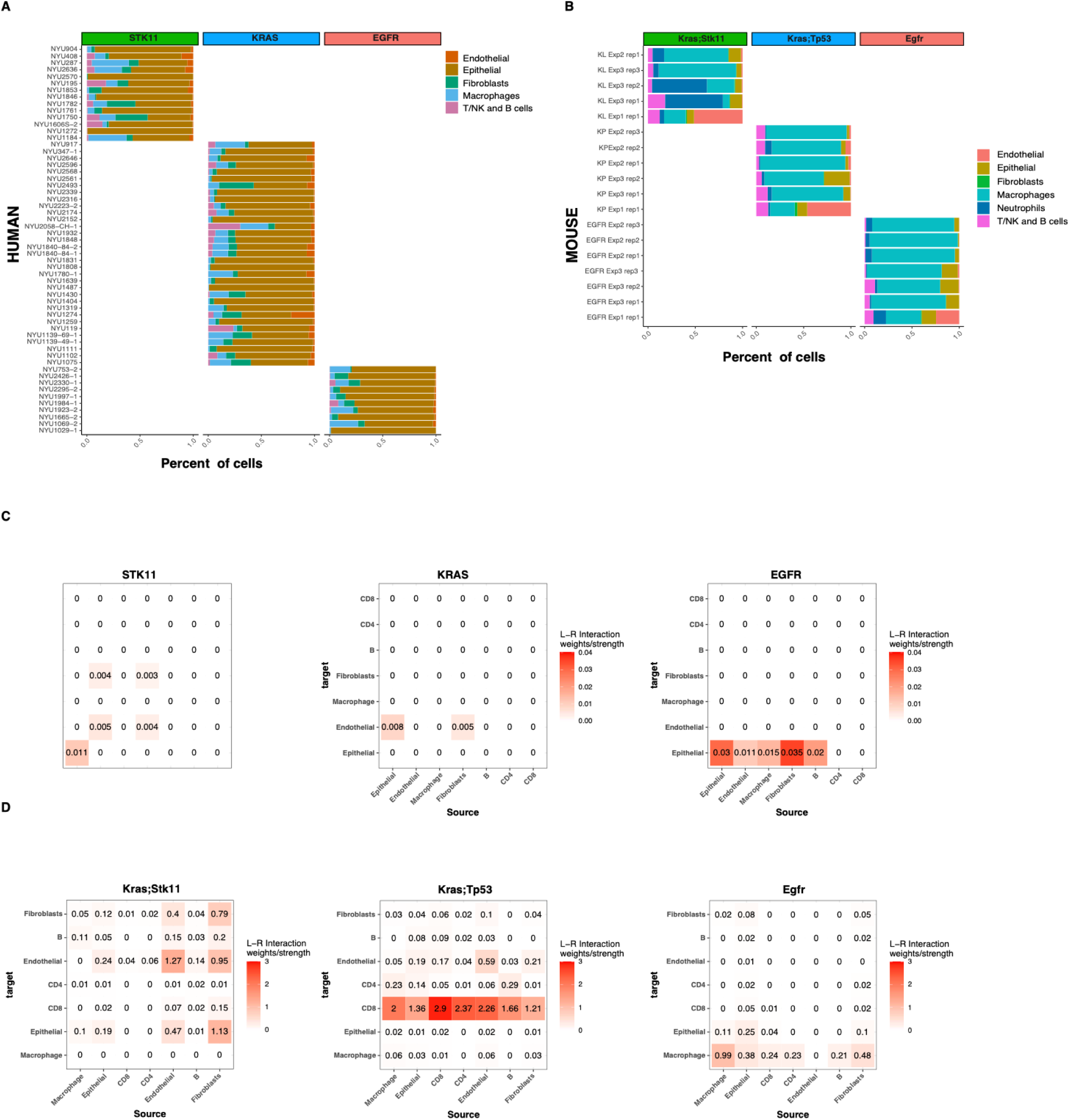
Cellular composition and communication of human and mouse LUAD samples. Stacked bar plots of cell-type percentages by individual (**A)** LUAD patients, (**B)** individual LUAD GEMMs. Correlation maps depicting the strength of cell interactions deriving from DEGs between genotype in **(C)** human and **(D)** mouse.

**Figure S3.**
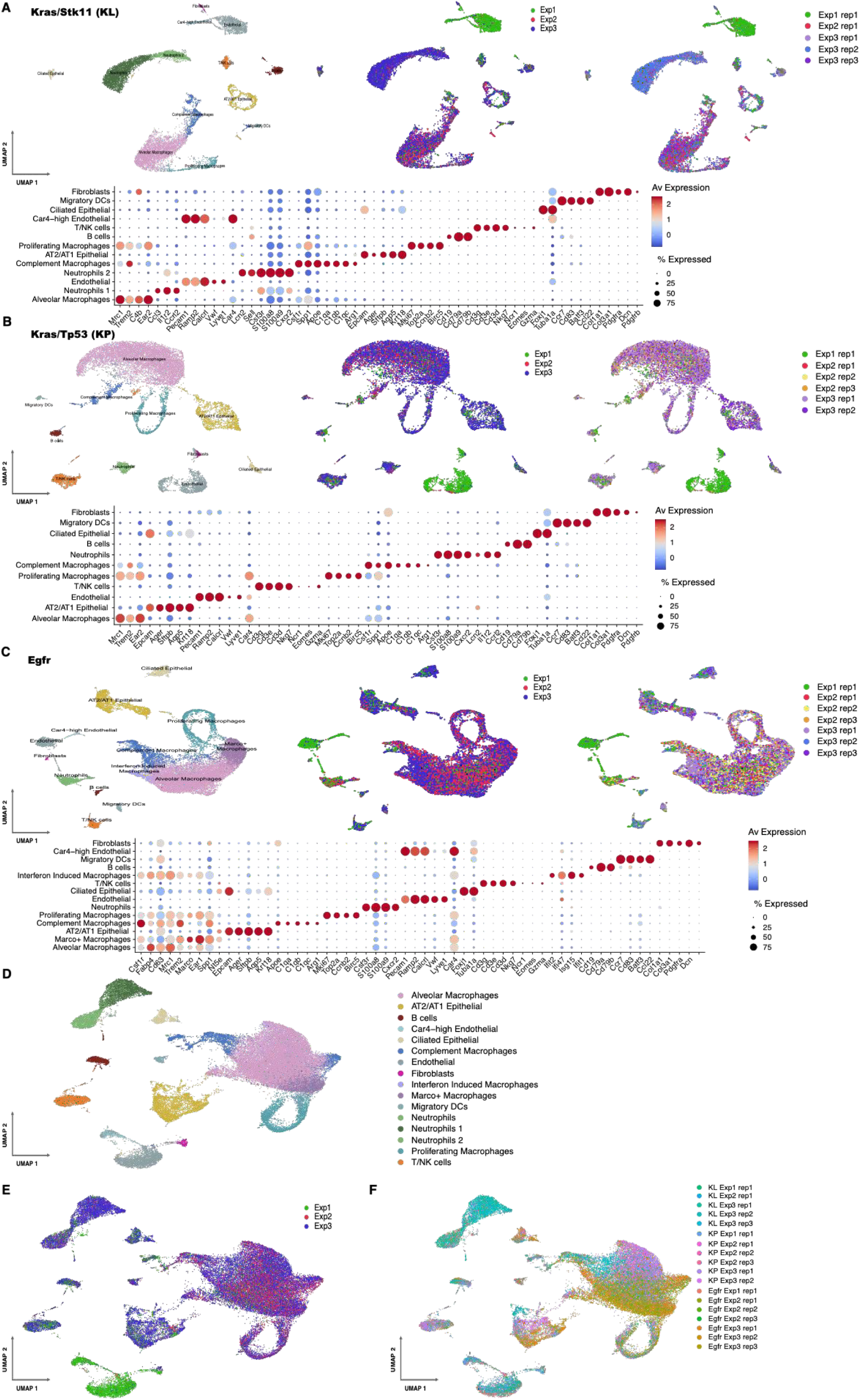
Single-cell RNA sequencing of lung adenocarcinoma GEMM specimens allows genotype-driven profiling. **A)** UMAP visualization of mouse *Kras/Stk11* (KL) LUAD specimens. Three independent experiments were performed and five mice were used in this study. Box plots indicate gene markers used for cell type identification. **B)** UMAP visualization of mouse *Kras/Tp53* (KP) LUAD specimens. Three independent experiments were performed, and six mice were used in this cohort. Box plots indicate gene markers used for cell type identification. **C)** UMAP visualization of mouse *Egfr* LUAD specimens. Three independent experiments were performed, and seven mice were used in this cohort. Box plots indicate gene markers used for cell type identification.**D–F)** UMAP visualization of integrated scRNA data from all 18 mice by cell type (**D)**, experiment (**E)**, and mouse (**F)**.

**Figure S4.**
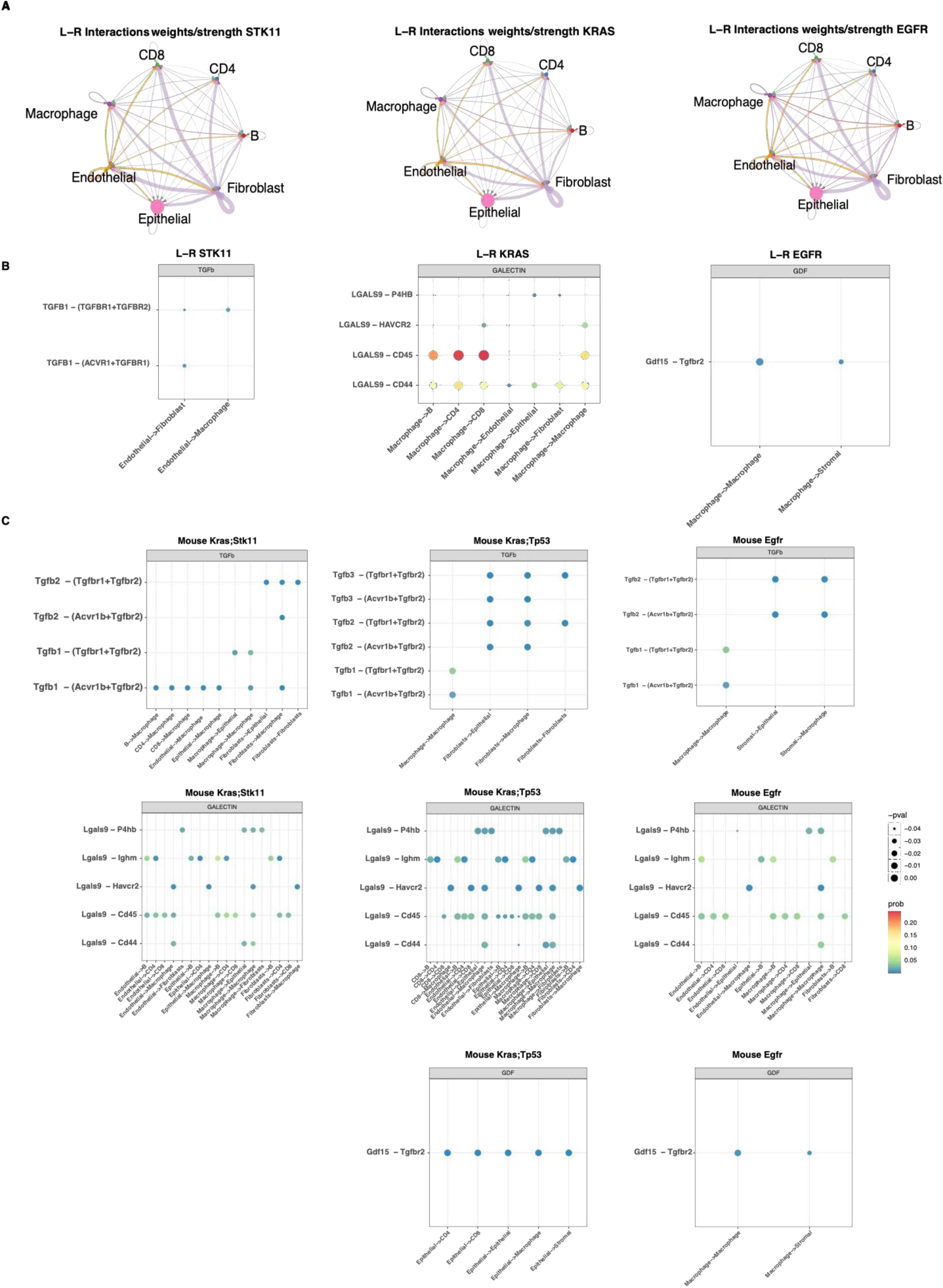
Genotype-stratified cell–cell communication networks in lung adenocarcinoma. **A)** Chord diagrams illustrating the landscape of intercellular interactions in human LUAD, stratified by oncogenic genotype. **B–C)** Genotype-specific interaction networks in human **(B)** and murine **(C)** LUAD, highlighting conserved and divergent communication patterns shaped by STK11, KRAS, and EGFR mutations.

**Figure S5.**
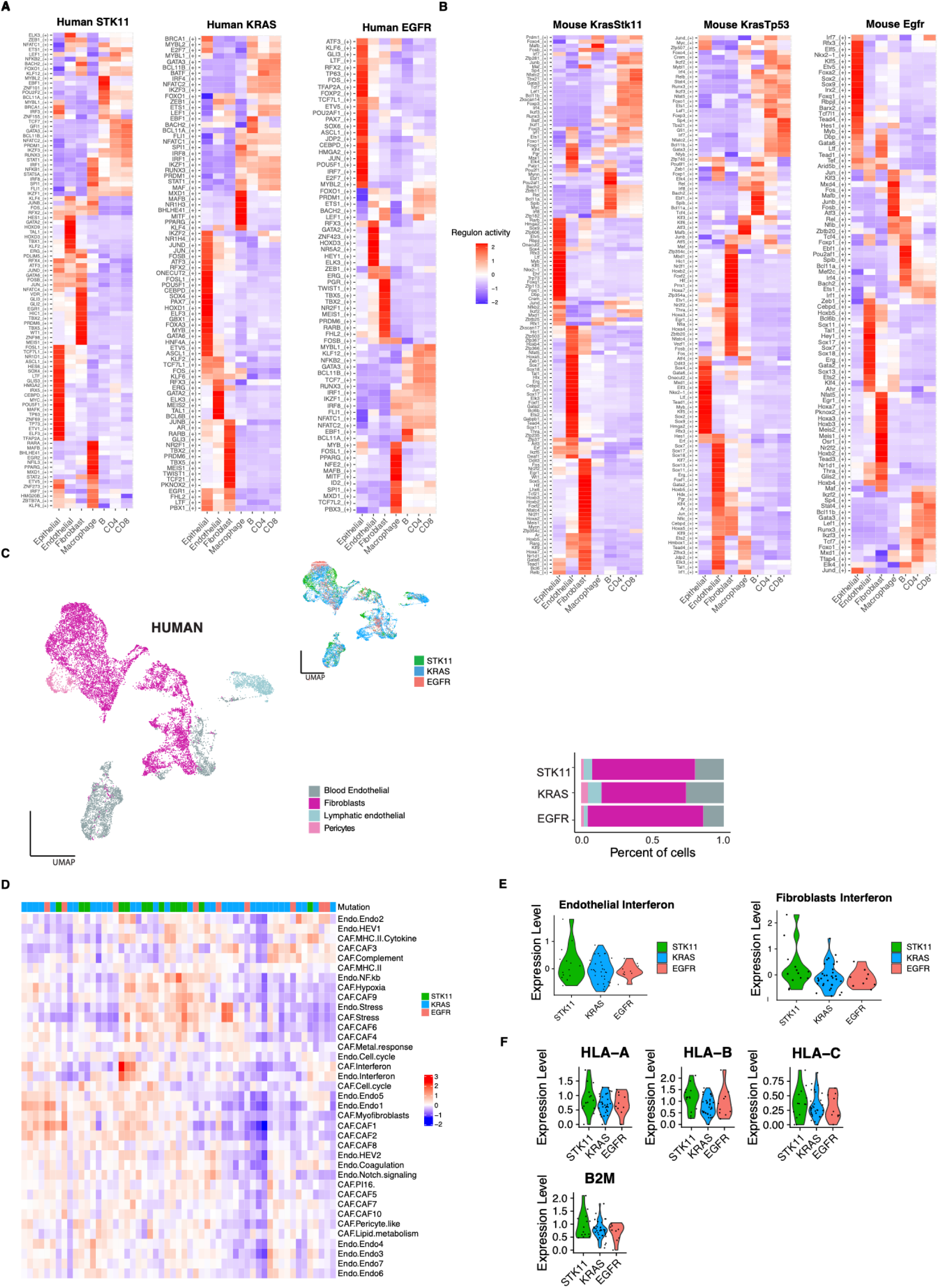
Transcriptional regulation and cellular programs across oncogenotypes in the LUAD tumor microenvironment. **A–B)** Heatmaps depicting the activity of regulons across STK11-, KRAS-, and EGFR-driven tumors in human **(A)** and mouse **(B)** LUAD specimens. **C)** UMAP projection and corresponding stacked bar plots illustrating the distribution of non-immune cell populations in the human LUAD microenvironment stratified by genotype. **D)** Heatmap displaying the enrichment scores of Gavish-defined modules across non-immune tumor microenvironment compartments in human LUAD. **E)** Violin plots showing differential expression of the interferon response module in endothelial and fibroblast populations across the three oncogenotypes in human LUAD. **F)** Expression levels of representative MHC class I complex genes across genotypes in the non-immune tumor microenvironment of human LUAD.

**Figure S6.**
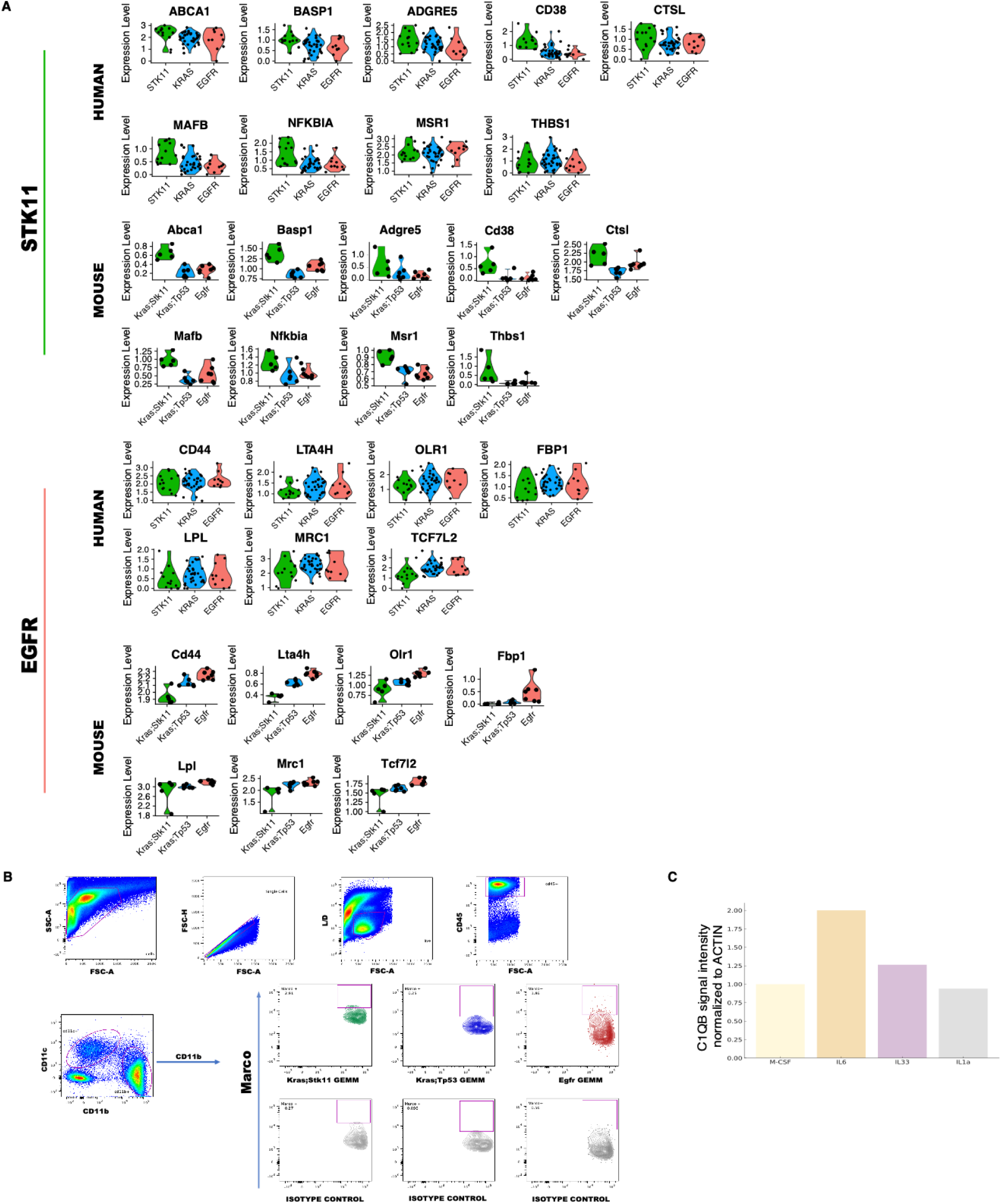
LUAD transcriptome analyses identify *MARCO* and *C1Q* as genotype-specific macrophage genes. **A)** Violin plots displaying the average per-patient and per-mouse expression of the identified cross-species shared genes in human LUAD macrophages. **B)** Flow cytometry gating strategy used to assess MARCO expression in TAMs of LUAD GEMMs and representative plots of MARCO expression in CD11b + cells. **C)** Barplots indicate the normalized signal intensity quantification of the C1QB/ACTIN ratio of the immunoblot in Fig. 4Q.

**Figure S7.**
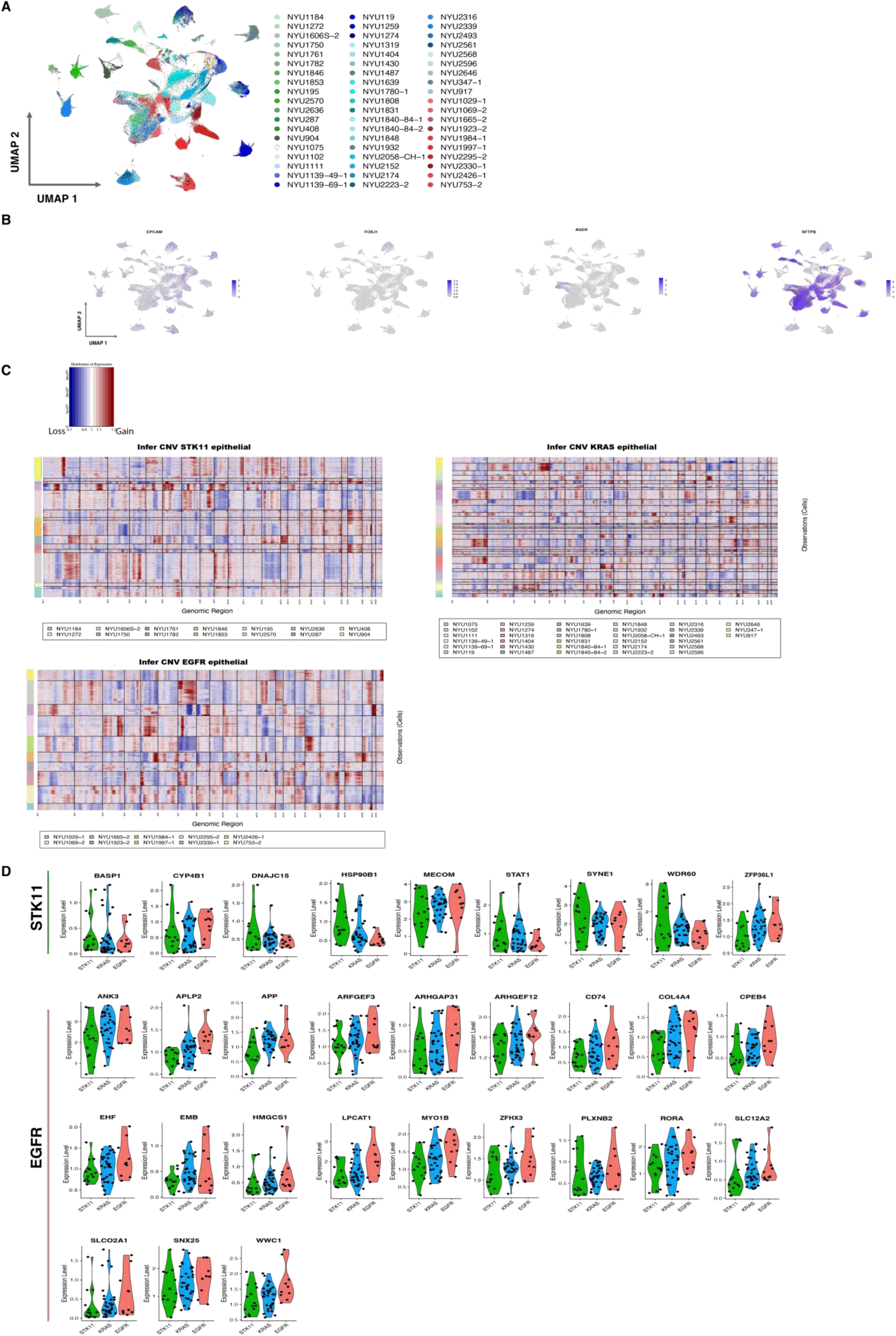
Copy number variation (CNV) analyses in human LUAD epithelial cells. **A)** UMAP visualization of epithelial cells from 57 human LUAD specimens using patient ID. **B)** UMAP projection of human epithelial markers. **C)** InferCNV (copy number variation) analyses in human LUAD epithelial cells verify the presence of large chromosomal copy number variations, such as gains (depicted in red) or deletions (depicted in blue). **D)** Violin plots displaying the average per-patient expression of the identified cross-species-shared genes in human LUAD epithelial cells.

**Figure S8.**
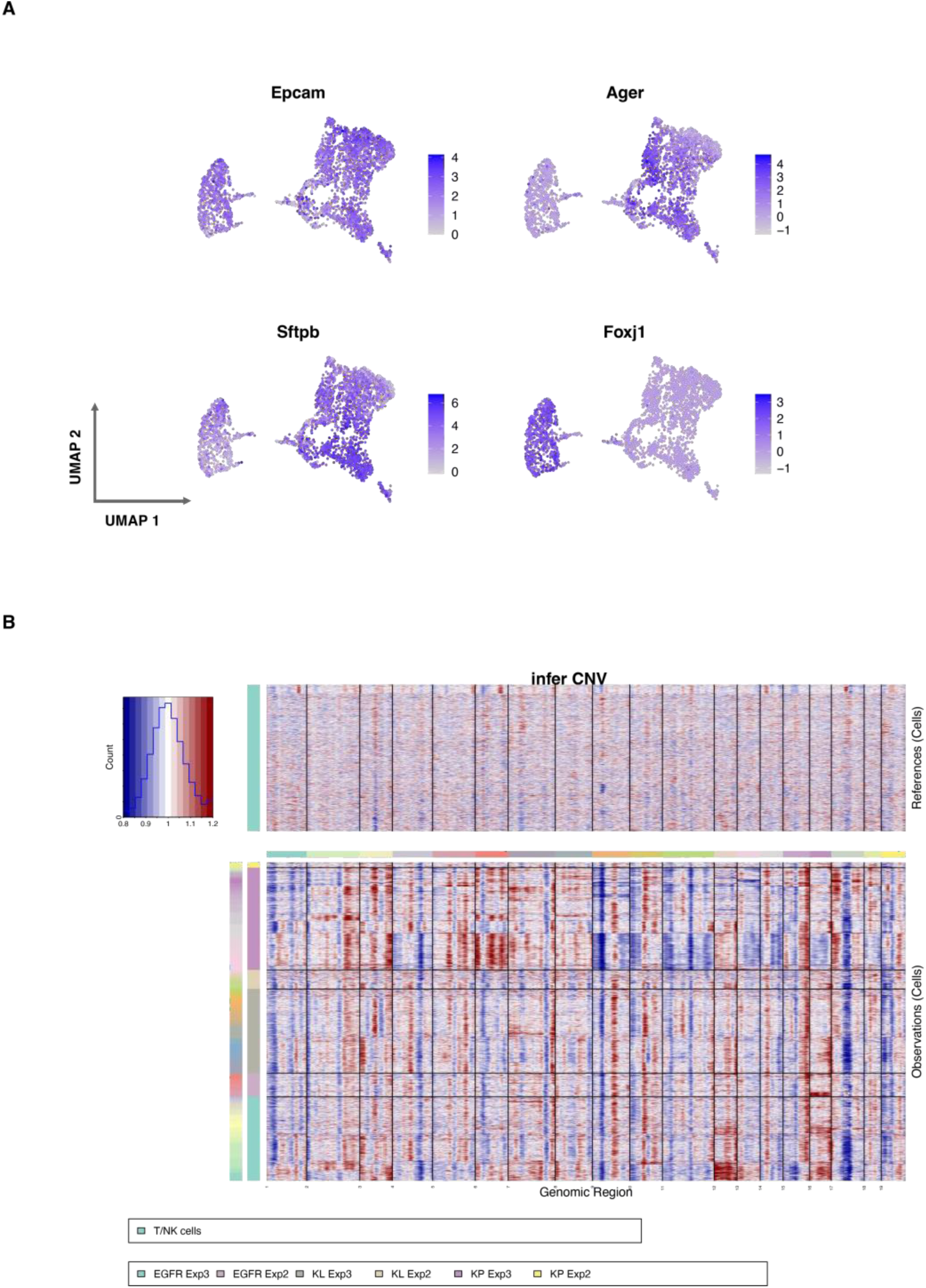
Copy number variation (CNV) analyses of epithelial cells from LUAD GEMMs. **A)** UMAP projection of mouse lung epithelial markers. **B)** InferCNV (copy number variation) analyses in mouse LUAD epithelial cells verify the presence of large chromosomal copy number variations, such as gains (depicted in red) or deletions (depicted in blue).

**Figure S9.**
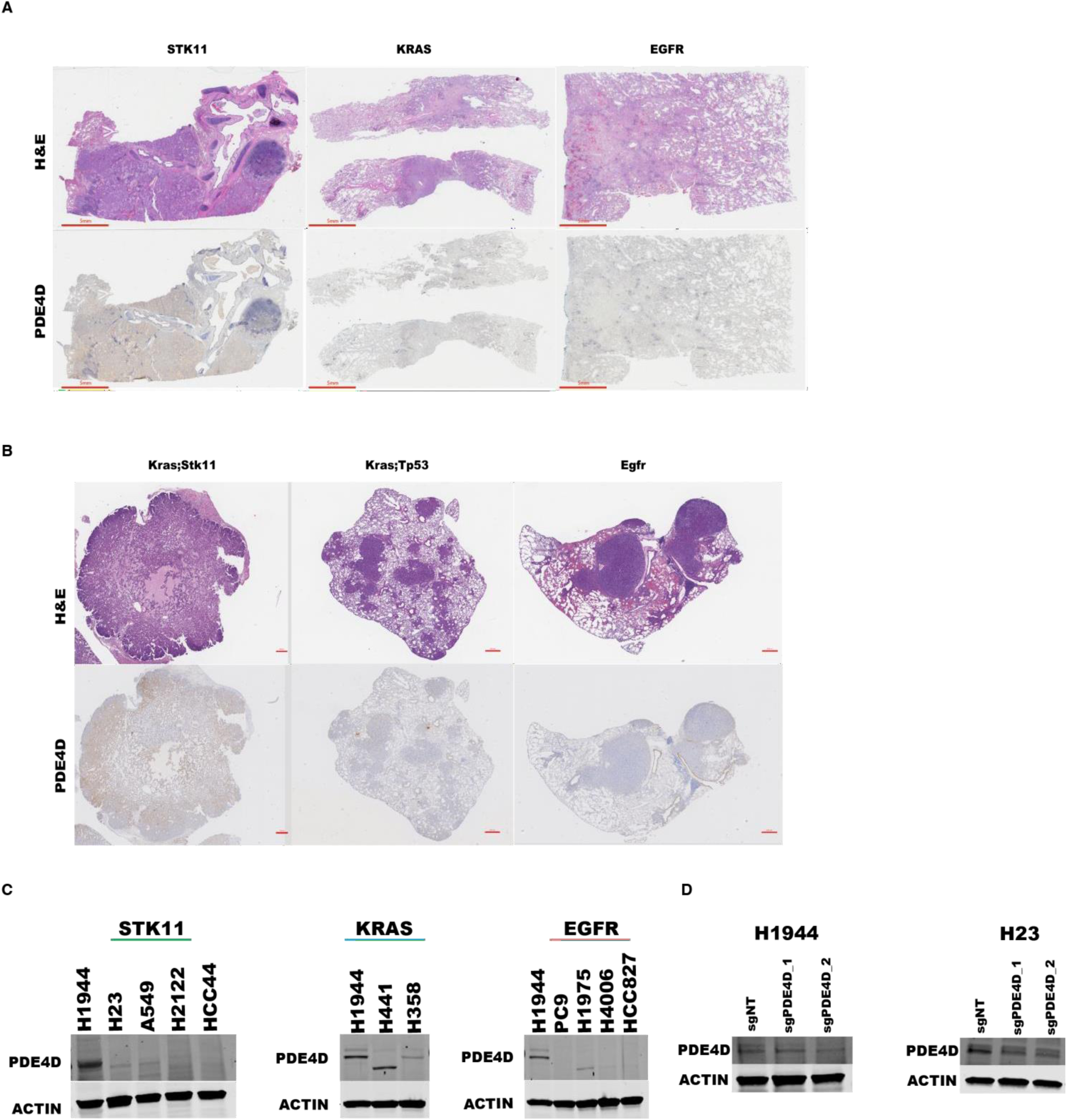
*PDE4D* is an *STK11* dependency in lung adenocarcinoma. **A)** Representative H&E and PDE4D staining images of patients with LUAD. Scale bars, 5 mm. **B)** Representative H&E and PDE4D staining of LUAD GEMMs. Scale bars, 100 μm. **C)** Immunoblots assessing the PDE4D expression levels in LUAD human cell lines. **D)** Immunoblots verifying CRISPR-mediated *PDE4D* ablation in the human H1944 and H23 LUAD cell lines. sgNT served as a non-targeting gRNA control. Actin served as a control to verify equal protein loading.

**Figure S10.**
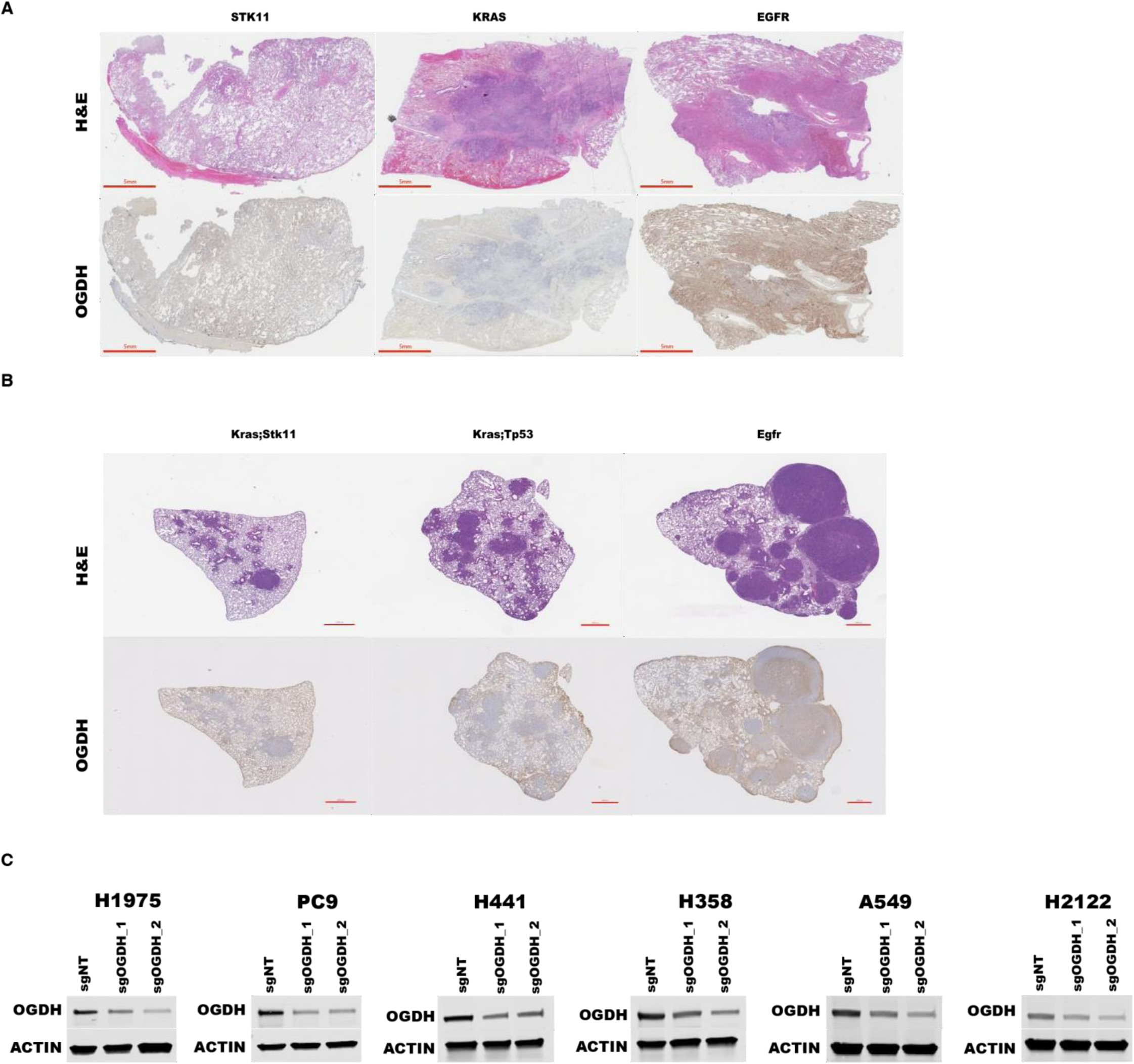
*OGDH* is a metabolic dependency in *EGFR*-mutated LUAD. **A)** Representative H&E and OGDH staining in patients with LUAD. Scale bars, 5 mm. **B)** Representative H&E and OGDH staining of LUAD GEMMs. Scale bars, 100 μm. **C)** Immunoblots verifying CRISPR-mediated *OGDH* ablation in human LUAD cell lines. sgNT served as a non-targeting gRNA control. Actin served as a control to verify equal protein loading.

## References

1. Siegel RL, Miller KD, Fuchs HE, Jemal A. Cancer statistics, 2022. CA Cancer J Clin 2022;72(1):7–33 doi 10.3322/caac.21708.

2. Leiter A, Veluswamy RR, Wisnivesky JP. The global burden of lung cancer: current status and future trends. Nat Rev Clin Oncol 2023;20(9):624–39 doi 10.1038/s41571-023-00798-3.

3. Molina JR, Yang P, Cassivi SD, Schild SE, Adjei AA. Non-small cell lung cancer: epidemiology, risk factors, treatment, and survivorship. Mayo Clin Proc 2008;83(5):584–94 doi 10.4065/83.5.584.

4. Passaro A, Al Bakir M, Hamilton EG, Diehn M, Andre F, Roy-Chowdhuri S, et al. Cancer biomarkers: Emerging trends and clinical implications for personalized treatment. Cell 2024;187(7):1617–35 doi 10.1016/j.cell.2024.02.041.

5. Cohen P, Cross D, Janne PA. Kinase drug discovery 20 years after imatinib: progress and future directions. Nat Rev Drug Discov 2021;20(7):551–69 doi 10.1038/s41573-021-00195-4.

6. Huang J, Pan H, Sun J, Wu J, Xuan Q, Wang J, et al. TMEM147 aggravates the progression of HCC by modulating cholesterol homeostasis, suppressing ferroptosis, and promoting the M2 polarization of tumor-associated macrophages. J Exp Clin Cancer Res 2023;42(1):286 doi 10.1186/s13046-023-02865-0.

7. Molina-Arcas M, Downward J. Exploiting the therapeutic implications of KRAS inhibition on tumor immunity. Cancer Cell 2024;42(3):338–57 doi 10.1016/j.ccell.2024.02.012.

8. Walsh RJ, Sundar R, Lim JSJ. Immune checkpoint inhibitor combinations-current and emerging strategies. Br J Cancer 2023;128(8):1415–7 doi 10.1038/s41416-023-02181-6.

9. Mountzios G, Remon J, Hendriks LEL, Garcia-Campelo R, Rolfo C, Van Schil P, et al. Immune-checkpoint inhibition for resectable non-small-cell lung cancer - opportunities and challenges. Nat Rev Clin Oncol 2023;20(10):664–77 doi 10.1038/s41571-023-00794-7.

10. Majem B, Nadal E, Munoz-Pinedo C. Exploiting metabolic vulnerabilities of Non small cell lung carcinoma. Semin Cell Dev Biol 2020;98:54–62 doi 10.1016/j.semcdb.2019.06.004.

11. Lavin Y, Kobayashi S, Leader A, Amir E-AD, Elefant N, Bigenwald C, et al. Innate Immune Landscape in Early Lung Adenocarcinoma by Paired Single-Cell Analyses. Cell 2017;169(4):750–65.e17 doi 10.1016/j.cell.2017.04.014.

12. Maynard A, McCoach CE, Rotow JK, Harris L, Haderk F, Kerr DL, et al. Therapy-Induced Evolution of Human Lung Cancer Revealed by Single-Cell RNA Sequencing. Cell 2020;182(5):1232–51 e22 doi 10.1016/j.cell.2020.07.017.

13. Koyama S, Akbay EA, Li YY, Aref AR, Skoulidis F, Herter-Sprie GS, et al. STK11/LKB1 Deficiency Promotes Neutrophil Recruitment and Proinflammatory Cytokine Production to Suppress T-cell Activity in the Lung Tumor Microenvironment. Cancer Res 2016;76(5):999–1008 doi 10.1158/0008-5472.CAN-15-1439.

14. Biton J, Mansuet-Lupo A, Pecuchet N, Alifano M, Ouakrim H, Arrondeau J, et al. TP53, STK11, and EGFR Mutations Predict Tumor Immune Profile and the Response to Anti-PD-1 in Lung Adenocarcinoma. Clin Cancer Res 2018;24(22):5710–23 doi 10.1158/1078-0432.CCR-18-0163.

15. Skoulidis F, Goldberg ME, Greenawalt DM, Hellmann MD, Awad MM, Gainor JF, et al. STK11/LKB1 Mutations and PD-1 Inhibitor Resistance in KRAS-Mutant Lung Adenocarcinoma. Cancer Discov 2018;8(7):822–35 doi 10.1158/2159-8290.CD-18-0099.

16. Hayes SA, Hudson AL, Clarke SJ, Molloy MP, Howell VM. From mice to men: GEMMs as trial patients for new NSCLC therapies. Semin Cell Dev Biol 2014;27:118–27 doi 10.1016/j.semcdb.2014.04.002.

17. McFadden DG, Politi K, Bhutkar A, Chen FK, Song X, Pirun M, et al. Mutational landscape of EGFR-, MYC-, and Kras-driven genetically engineered mouse models of lung adenocarcinoma. Proc Natl Acad Sci U S A 2016;113(42):E6409–E17 doi 10.1073/pnas.1613601113.

18. Kim N, Kang H, Jo A, Yoo SA, Lee HO. Perspectives on single-nucleus RNA sequencing in different cell types and tissues. J Pathol Transl Med 2023;57(1):52–9 doi 10.4132/jptm.2022.12.19.

19. Skoulidis F, Byers LA, Diao L, Papadimitrakopoulou VA, Tong P, Izzo J, et al. Co-occurring genomic alterations define major subsets of KRAS-mutant lung adenocarcinoma with distinct biology, immune profiles, and therapeutic vulnerabilities. Cancer Discov 2015;5(8):860–77 doi 10.1158/2159-8290.CD-14-1236.

20. Deng J, Peng DH, Fenyo D, Yuan H, Lopez A, Levin DS, et al. In vivo metabolomics identifies CD38 as an emergent vulnerability in LKB1 -mutant lung cancer. bioRxiv 2023 doi 10.1101/2023.04.18.537350.

21. Wohlhieter CA, Richards AL, Uddin F, Hulton CH, Quintanal-Villalonga A, Martin A, et al. Concurrent Mutations in STK11 and KEAP1 Promote Ferroptosis Protection and SCD1 Dependence in Lung Cancer. Cell Rep 2020;33(9):108444 doi 10.1016/j.celrep.2020.108444.

22. Sutton P, Borgia JA, Bonomi P, Plate JM. Lyn, a Src family kinase, regulates activation of epidermal growth factor receptors in lung adenocarcinoma cells. Mol Cancer 2013;12:76 doi 10.1186/1476-4598-12-76.

23. Yue F, Jiang W, Ku AT, Young AIJ, Zhang W, Souto EP, et al. A Wnt-Independent LGR4-EGFR Signaling Axis in Cancer Metastasis. Cancer Res 2021;81(17):4441–54 doi 10.1158/0008-5472.CAN-21-1112.

24. Deng W, Gu L, Li X, Zheng J, Zhang Y, Duan B, et al. CD24 associates with EGFR and supports EGF/EGFR signaling via RhoA in gastric cancer cells. J Transl Med 2016;14:32 doi 10.1186/s12967-016-0787-y.

25. Yang R, Sun L, Li CF, Wang YH, Yao J, Li H, et al. Galectin-9 interacts with PD-1 and TIM-3 to regulate T cell death and is a target for cancer immunotherapy. Nat Commun 2021;12(1):832 doi 10.1038/s41467-021-21099-2.

26. Patel S, Alvarez-Guaita A, Melvin A, Rimmington D, Dattilo A, Miedzybrodzka EL, et al. GDF15 Provides an Endocrine Signal of Nutritional Stress in Mice and Humans. Cell Metab 2019;29(3):707–18 e8 doi 10.1016/j.cmet.2018.12.016.

27. Szabo PA, Levitin HM, Miron M, Snyder ME, Senda T, Yuan J, et al. Single-cell transcriptomics of human T cells reveals tissue and activation signatures in health and disease. Nat Commun 2019;10(1):4706 doi 10.1038/s41467-019-12464-3.

28. Gavish A, Tyler M, Greenwald AC, Hoefflin R, Simkin D, Tschernichovsky R, et al. Hallmarks of transcriptional intratumour heterogeneity across a thousand tumours. Nature 2023;618(7965):598-606 doi 10.1038/s41586-023-06130-4.

29. McEwan DG, Brunton VG, Baillie GS, Leslie NR, Houslay MD, Frame MC. Chemoresistant KM12C colon cancer cells are addicted to low cyclic AMP levels in a phosphodiesterase 4-regulated compartment via effects on phosphoinositide 3-kinase. Cancer Res 2007;67(11):5248–57 doi 10.1158/0008-5472.CAN-07-0097.

30. Rahrmann EP, Collier LS, Knutson TP, Doyal ME, Kuslak SL, Green LE, et al. Identification of PDE4D as a proliferation promoting factor in prostate cancer using a Sleeping Beauty transposon-based somatic mutagenesis screen. Cancer Res 2009;69(10):4388–97 doi 10.1158/0008-5472.CAN-08-3901.

31. Delyon J, Servy A, Laugier F, Andre J, Ortonne N, Battistella M, et al. PDE4D promotes FAK-mediated cell invasion in BRAF-mutated melanoma. Oncogene 2017;36(23):3252–62 doi 10.1038/onc.2016.469.

32. Kashima Y, Shibahara D, Suzuki A, Muto K, Kobayashi IS, Plotnick D, et al. Single-Cell Analyses Reveal Diverse Mechanisms of Resistance to EGFR Tyrosine Kinase Inhibitors in Lung Cancer. Cancer Res 2021;81(18):4835–48 doi 10.1158/0008-5472.CAN-20-2811.

33. Ding J, Ding X, Leng Z. LPCAT1 promotes gefitinib resistance via upregulation of the EGFR/PI3K/AKT signaling pathway in lung adenocarcinoma. J Cancer 2022;13(6):1837–47 doi 10.7150/jca.66126.

34. Bi J, Ichu TA, Zanca C, Yang H, Zhang W, Gu Y, et al. Oncogene Amplification in Growth Factor Signaling Pathways Renders Cancers Dependent on Membrane Lipid Remodeling. Cell Metab 2019;30(3):525–38 e8 doi 10.1016/j.cmet.2019.06.014.

35. Millman SE, Chaves-Perez A, Janaki-Raman S, Ho YJ, Morris JPt, Narendra V, et al. alpha-Ketoglutarate dehydrogenase is a therapeutic vulnerability in acute myeloid leukemia. Blood 2025;145(13):1422–36 doi 10.1182/blood.2024025245.

36. Nguyen TT, Torrini C, Shang E, Shu C, Mun JY, Gao Q, et al. OGDH and Bcl-xL loss causes synthetic lethality in glioblastoma. JCI Insight 2024;9(8) doi 10.1172/jci.insight.172565.

37. Lu X, Yang P, Zhao X, Jiang M, Hu S, Ouyang Y, et al. OGDH mediates the inhibition of SIRT5 on cell proliferation and migration of gastric cancer. Exp Cell Res 2019;382(2):111483 doi 10.1016/j.yexcr.2019.06.028.

38. Afshar-Kharghan V. The role of the complement system in cancer. J Clin Invest 2017;127(3):780–9 doi 10.1172/JCI90962.

39. Reis ES, Mastellos DC, Ricklin D, Mantovani A, Lambris JD. Complement in cancer: untangling an intricate relationship. Nat Rev Immunol 2018;18(1):5–18 doi 10.1038/nri.2017.97.

40. Bulla R, Tripodo C, Rami D, Ling GS, Agostinis C, Guarnotta C, et al. C1q acts in the tumour microenvironment as a cancer-promoting factor independently of complement activation. Nat Commun 2016;7:10346 doi 10.1038/ncomms10346.

41. Roumenina LT, Daugan MV, Noe R, Petitprez F, Vano YA, Sanchez-Salas R, et al. Tumor Cells Hijack Macrophage-Produced Complement C1q to Promote Tumor Growth. Cancer Immunol Res 2019;7(7):1091–105 doi 10.1158/2326-6066.CIR-18-0891.

42. Koyama S, Akbay EA, Li YY, Aref AR, Skoulidis F, Herter-Sprie GS, et al. STK11/LKB1 Deficiency Promotes Neutrophil Recruitment and Proinflammatory Cytokine Production to Suppress T-cell Activity in the Lung Tumor Microenvironment. Cancer Research 2016;76(5):999–1008 doi 10.1158/0008-5472.can-15-1439.

43. Kou W, Li B, Shi Y, Zhao Y, Yu Q, Zhuang J, et al. High complement protein C1q levels in pulmonary fibrosis and non-small cell lung cancer associated with poor prognosis. BMC Cancer 2022;22(1):110 doi 10.1186/s12885-021-08912-3.

44. Mastellos DC, Ricklin D, Lambris JD. Clinical promise of next-generation complement therapeutics. Nat Rev Drug Discov 2019;18(9):707–29 doi 10.1038/s41573-019-0031-6.

45. Sun Y, Wirta D, Murahashi W, Mathur V, Sankaranarayanan S, Taylor LK, et al. Safety and Target Engagement of Complement C1q Inhibitor ANX007 in Neurodegenerative Eye Disease: Results from Phase I Studies in Glaucoma. Ophthalmol Sci 2023;3(2):100290 doi 10.1016/j.xops.2023.100290.

46. Georgoudaki AM, Prokopec KE, Boura VF, Hellqvist E, Sohn S, Ostling J, et al. Reprogramming Tumor-Associated Macrophages by Antibody Targeting Inhibits Cancer Progression and Metastasis. Cell Rep 2016;15(9):2000–11 doi 10.1016/j.celrep.2016.04.084.

47. La Fleur L, Botling J, He F, Pelicano C, Zhou C, He C, et al. Targeting MARCO and IL37R on Immunosuppressive Macrophages in Lung Cancer Blocks Regulatory T Cells and Supports Cytotoxic Lymphocyte Function. Cancer Res 2021;81(4):956–67 doi 10.1158/0008-5472.CAN-20-1885.

48. Eisinger S, Sarhan D, Boura VF, Ibarlucea-Benitez I, Tyystjarvi S, Oliynyk G, et al. Targeting a scavenger receptor on tumor-associated macrophages activates tumor cell killing by natural killer cells. Proc Natl Acad Sci U S A 2020;117(50):32005–16 doi 10.1073/pnas.2015343117.

49. Sa JK, Chang N, Lee HW, Cho HJ, Ceccarelli M, Cerulo L, et al. Transcriptional regulatory networks of tumor-associated macrophages that drive malignancy in mesenchymal glioblastoma. Genome Biol 2020;21(1):216 doi 10.1186/s13059-020-02140-x.

50. Ferrarone JR, Thomas J, Unni AM, Zheng Y, Nagiec MJ, Gardner EE, et al. Genome-wide CRISPR screens in spheroid culture reveal that the tumor suppressor LKB1 inhibits growth via the PIKFYVE lipid kinase. Proc Natl Acad Sci U S A 2024;121(21):e2403685121 doi 10.1073/pnas.2403685121.

51. Yin D, Lu X, Liang X, Lu Y, Xiong L, Wu P, et al. STK11 genetic alterations in metastatic EGFR mutant lung cancer. Sci Rep 2025;15(1):5729 doi 10.1038/s41598-024-74779-6.

52. Cheng FJ, Chen CH, Tsai WC, Wang BW, Yu MC, Hsia TC, et al. Cigarette smoke-induced LKB1/AMPK pathway deficiency reduces EGFR TKI sensitivity in NSCLC. Oncogene 2021;40(6):1162–75 doi 10.1038/s41388-020-01597-1.

53. Sakkas LI, Mavropoulos A, Bogdanos DP. Phosphodiesterase 4 Inhibitors in Immune-mediated Diseases: Mode of Action, Clinical Applications, Current and Future Perspectives. Curr Med Chem 2017;24(28):3054–67 doi 10.2174/0929867324666170530093902.

54. Schafer PH, Parton A, Gandhi AK, Capone L, Adams M, Wu L, et al. Apremilast, a cAMP phosphodiesterase-4 inhibitor, demonstrates anti-inflammatory activity in vitro and in a model of psoriasis. Br J Pharmacol 2010;159(4):842–55 doi 10.1111/j.1476-5381.2009.00559.x.

55. Kaminow B, Yunusov D, Dobin A. 2021 doi 10.1101/2021.05.05.442755.

56. 56. Berg M, Petoukhov I, van den Ende I, Meyer KB, Guryev V, Vonk JM, et al. FastCAR: fast correction for ambient RNA to facilitate differential gene expression analysis in single-cell RNA-sequencing datasets. BMC Genomics 2023;24(1):722 doi 10.1186/s12864-023-09822-3.

57. Bernstein NJ, Fong NL, Lam I, Roy MA, Hendrickson DG, Kelley DR. Solo: Doublet Identification in Single-Cell RNA-Seq via Semi-Supervised Deep Learning. Cell Syst 2020;11(1):95–101 e5 doi 10.1016/j.cels.2020.05.010.

58. Gayoso A, Lopez R, Xing G, Boyeau P, Valiollah Pour Amiri V, Hong J, et al. A Python library for probabilistic analysis of single-cell omics data. Nat Biotechnol 2022;40(2):163–6 doi 10.1038/s41587-021-01206-w.

59. Hafemeister C, Satija R. Normalization and variance stabilization of single-cell RNA-seq data using regularized negative binomial regression. Genome Biol 2019;20(1):296 doi 10.1186/s13059-019-1874-1.

